# A Markov constraint to uniquely identify elementary flux mode weights in unimolecular metabolic networks

**DOI:** 10.1101/2022.07.25.501464

**Authors:** Justin G. Chitpin, Theodore J. Perkins

## Abstract

Elementary flux modes (EFMs) are minimal, steady state pathways characterizing a flux network. Fundamentally, all steady state fluxes in a network are decomposable into a linear combination of EFMs. While there is typically no unique set of EFM weights that reconstructs these fluxes, several optimization-based methods have been proposed to constrain the solution space by enforcing some notion of parsimony. However, it has long been recognized that optimization-based approaches may fail to uniquely identify EFM weights and return different feasible solutions across objective functions and solvers. Here we show that, for flux networks only involving single molecule transformations, these problems can be avoided by imposing a Markovian constraint on EFM weights. Our Markovian constraint guarantees a unique solution to the flux decomposition problem, and that solution is arguably more biophysically plausible than other solutions. We describe an algorithm for computing Markovian EFM weights via steady state analysis of a certain discrete-time Markov chain, based on the flux network, which we call the cycle-history Markov chain. We demonstrate our method with a differential analysis of EFM activity in a lipid metabolic network comparing healthy and Alzheimer’s disease patients. Our method is the first to uniquely decompose steady state fluxes into EFM weights for any unimolecular metabolic network.

## 1 Introduction

Cellular metabolism consists of carefully-tuned biochemical reactions operating collectively to maintain metabolic homeostasis. These reactions form highly connected metabolic networks whose dynamics regulate bioenergetic processes [1, 2] and cell-specific functions [3]. Data from large-scale genomics projects has led to genome-wide reconstructions of bacterial, yeast, plant, and human metabolic networks [4–7].

When metabolic reaction rates are known, the relationships between metabolites can be modelled as a flux network with metabolites represented as nodes and the fluxes between them as weighted, directed edges. The fluxes may be estimated experimentally by stable isotope tracing analysis and metabolic flux analysis [8, 9] or computationally by flux balance analysis [10], kinetic models [11, 12], and other statistical techniques [13, 14]. Computational methods generally estimate fluxes under metabolic steady state, where the production of each metabolite is balanced by its consumption. These flux networks are often constructed to visualize metabolic activity, or analyzed to optimize metabolite production [15] and identify metabolic alterations associated with diseases [16, 17].

Elementary flux modes (EFMs) are a pathway-based approach to study properties of flux networks. The EFM is defined as a non-decomposable, steady state pathway in a flux network [18]. EFMs may either traverse the network, connecting input and output fluxes of external metabolites, or encode looped pathways, such as futile cycles, to maintain steady state. The non-decomposable property specifies that pathway flux is lost if any reaction in an EFM is abolished. Thus, EFMs have been described as functional units of flux expressing any steady state metabolic phenotype [19].

EFM analyses have been extensively applied in literature to characterize metabolic activity. For example, the number of EFMs producing a given metabolite is an indicator of network robustness. Considerable work has been devoted to quantifying the contribution of individual EFMs towards external metabolite production of bioreactors [20] and correlating EFM activity with metabolic phenotypes [21]. More recently, EFM analyses predicted metabolic interactions in microbial communities [22], estimated cellular processes explaining exponential cell growth [23], and optimized renewable fuel production in bioreactors [24].

A major computational challenge associated with EFMs is their enumeration in large-scale networks. EFM analyses are typically limited to small biological networks (*<* 100 reactions) because the number of EFMs explodes combinatorially with network size and number of external metabolites [25, 26]. Considerable attention has been devoted to scaling EFM enumeration towards genome-wide networks [27–29]. Most recently, advances in parallelized EFM computation have enabled EFM enumeration in medium-sized networks containing as many as 296 reactions with over twelve billion EFMs [30].

Another key computational problem, and the focus of the current manuscript, is to determine how much flux is travelling along each EFM, given the steady state flux along each reaction in a metabolic network. This flux decomposition problem commonly arises in differential EFM analyses to identify metabolic subpathways altered across disease-versus-control or growth conditions [31, 32]. However, decomposing steady state fluxes onto EFMs is generally not unique. In a unimolecular flux network with *M* metabolites, the steady state phenotype can be described by the *N* reaction fluxes, of which there are at most *M* ^2^. In general, the number of EFMs, *K*, can be as large as *M* factorial, depending on the structure of the network.

Most existing strategies for solving the flux decomposition problem are dependent on some notion of parsimony and computational elegance, rather than strong biophysical motivation. These optimization-based methods use linear, quadratic, or mixed-integer linear programming (LP/QP/MILP) to assign EFM weights according to a user-specified objective function. Common optimization-based methods minimize or maximize activity through the shortest or fewest EFMs to reconstruct the observed fluxes. However, these methods may return one among many equally good solutions, often without acknowledging the other possibilities.

This issue of non-uniqueness is well-documented in the related metabolic field of flux balance analysis (FBA) which uses similar optimization-based techniques to predict steady state flux distributions [33, 34]. Many FBA objective functions have been proposed to quantify the contribution of each reaction towards the metabolic phenotype, producing different flux distributions [35] or a range of equally feasible flux distributions [36–38]. Advancements in FBA are solving the non-uniqueness problem by refining the objective function with additional “-omics” data to better describe the metabolic phenotype [39–42]. While this strategy may improve FBA flux predictions, it is unclear how transcript, metabolite, or protein levels could be leveraged to improve EFM weight identification.

Figure 1 highlights the flux decomposition problem on two subnetworks taken from the KEGG database [43]. In the first network, glutamate is catabolized via two parallel pathways resulting in proline synthesis. The second network shows THF remodelling into three derivatives within the one carbon folate cycle. Figure 1a shows five arbitrary sets of EFM weights that perfectly reconstruct the steady state fluxes. Applying an objective function to minimize the number of active EFMs would yield two equally good integer solutions, while a maximization function would return an infinite number of EFM weight solutions. Similar problems arise in Figure 1b, where different objective functions may either fail to constrain the solution space, or disagree on the unique set of EFM weights based on the chosen constraints. For example, prioritizing EFM activity through the shortest EFMs in Figure 1b would yield the first example solution, while optimizing for the fewest number or activity through the longest possible EFMs would return the second solution.

**Figure 1:**
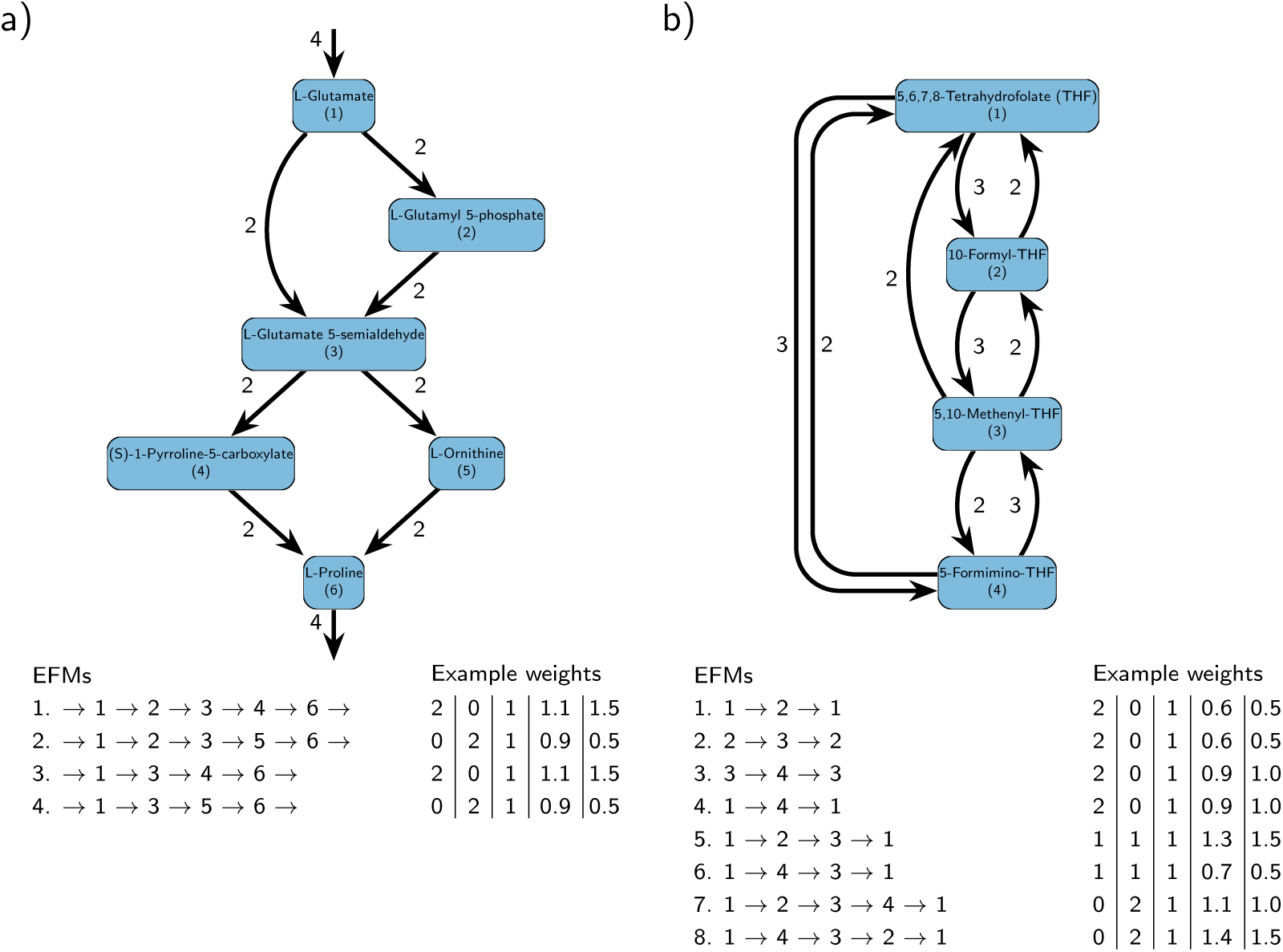
KEGG networks demonstrating EFM weight ambiguity. Nodes and edges denote metabolites and fluxes, respectively. EFMs weights that reconstruct the observed fluxes are shown for the highlighted nodes/edges. a) Pathway of L-Glutamate to L-Proline (KEGG rn00330). b) Subnetwork of one carbon folate cycle from (KEGG map00670).

When flux decompositions are not unique, the resulting EFM weights lose their metabolic predictivity. Fundamentally, the chosen EFM weights are a biophysical explanation of how substrates are metabolized in a network. Only a single set of EFM weights should correctly describe the metabolic processes governing the observed, steady state fluxes. We may verify this proposition by considering the set of EFM weights reconstructing the fluxes in Figure 1a. Decomposing the fluxes onto EFMs 1 and 4 would suggest glutamate is metabolized separately through two parallel paths that converge on proline. Deleting the reaction between L-glutamate and L-glutamate 5-semialdehyde should therefore abolish flux through (S)-1-pyrroline-5-carboxylate. Conversely, if the fluxes were instead explained by EFMs 2 and 3, we would expect a loss of flux through L-ornithine. This same problem occurs in Figure 1b where each chosen set of EFM weights offers a different biological explanation of THF remodelling.

Despite the well-established use of mathematical optimization, uniquely identifying EFM weights remains a longstanding problem limiting EFM applications. In this article, we show how common objective functions described in literature may fail to unambiguously decompose fluxes onto EFM weights. We further argue these methods can produce biophysically unreasonable solutions. A metabolite has no memory of the reactions leading to its current state nor the sequence of reactions it will undergo to complete an EFM in the future. We argue that a more natural approach to assigning EFM weights is to model how a single particle converts between metabolites according to the observed, steady state fluxes. This Markovian assumption captures the intuition of the uniform EFM weight solution best describing the steady state fluxes in Figure 1a, and motivates our discrete-time Markov chain model of a single particle converting between metabolites in a unimolecular flux network. By decomposing the Markov chain trajectories into EFMs, we compute a unique probability of a particle transiting an EFM that is proportional to the EFM weights reconstructing those observed fluxes. We solve the problem of computing the EFM transit probabilities via steady state analysis of a new construction we propose, a cycle-history Markov chain. We demonstrate our method by computing EFM weights for the example flux networks in Figure 1 and a kinetic model of sphingolipid metabolism of healthy versus Alzheimer’s disease patients.

## 2 Methods

### 2.1 Flux networks and elementary flux modes

A unimolecular, steady-state flux network can be represented as a weighted graph *G* = (*V, E, f*), where *V* is the set of nodes corresponding to distinct metabolites, *E* is the set of directed edges corresponding to chemical transformations of one metabolite into another, and *f* are positive real-valued weights of those edges that denote the amount of metabolite per unit time being transformed [44]. This network may be closed or open such that external fluxes enter and leave via source and sink metabolites. In an open network, we assume there is a special node *v_ext_ ∈ V* (or possibly multiple such nodes) that represents the external environment. Flux into the network happens along edges *e* = (*v_ext_, i*) *∈ E* for which *i ∈ V −{v_ext_}* and *f* (*e*) *>* 0. Such nodes *i* are called sources. Flux out of the network happens along edges *e* = (*i, v_ext_*) *∈ E* for which *v ∈ V − {v_ext_}* and *f* (*e*) *>* 0, with such nodes *i* being called sinks. Note that open networks with sources and sinks can be modelled in other ways. However, this way is most notationally convenient for our purposes, because it allows open and closed networks to be treated simultaneously.

We restrict attention to flux networks at *steady state*, meaning that the total flux into each node equals the total flux out: for every *i ∈ V*, ∑*_j̸_*_=_*_i_ f_ji_* = *_j̸_*_=_*_i_ f_ij_*. We also restrict attention to *strongly connected* flux networks, where there must be a path in the network from any node *i* to any other node *j*. For open networks, this includes the possibility of paths through the external node(s) *v_ext_*. If one wants to analyze a flux network with multiple strongly connected components, then each component can be analyzed independently by our proposed method.

The flux network *G* can be decomposed into a finite set of *K* EFMs transiting a series of edges in *E* [45]. In the present context, the EFMs are the set of all simple cycles in *G*. The steady state fluxes in *G* can always be explained as a positive, linear combination of EFMs [46]. Specifically, the weighted sum of all EFMs contributing towards the same reaction should equal the observed, steady state flux of that reaction. This flux decomposition problem can be solved by identifying a set of EFM weights *w* satisfying *Aw* = *f*, where *A* is an *|E|* by *K* matrix with one or zero elements denoting whether a unimolecular reaction is active or inactive in each EFM. When the number of EFMs exceeds the number of linearly independent fluxes, *w* is undetermined. While optimization-based methods are not guaranteed to constrain the solution space of *w*, the method we present uniquely identifies *w*.

### 2.2 Markovian solution to the flux decomposition problem

We propose *Markovian EFM weights* as a way of resolving the issue that finding EFM weights to explain a given set of fluxes is generally an underdetermined problem. We develop our approach in three main steps. First, we define a Markov chain that models a “particle” of metabolite flowing randomly around the flux network and the steady state frequencies with which that particle passes around different simple cycles (i.e. EFMs). Second, we propose a novel construction, the *cycle-history Markov chain*, to efficiently calculate those frequencies. Third and finally, we describe how to scale those frequencies to calculate a unique Markovian solution to the EFM weight problem. While we provide intuitive justification of these steps, Supplementary information section 1 contains a complete proof of our algorithm correctness.

#### 2.2.1 Single particle, discrete-time Markov chain model

Our approach to uniquely identifying EFM weights depends on imagining a single “particle” transiting the flux network–repeatedly transformed from one metabolite to another. We make a Markovian assumption that when the particle is in metabolite state *i ∈ V*, it transitions to a new metabolite state *j* with probabilities proportional to the outgoing fluxes *f_ij_*. This assumption is very natural from a biophysical standpoint. If metabolite molecules are well mixed in a reaction volume along with transforming enzymes, then the future behaviour of any molecule should depend only on its current state. This situation is modelled by a discrete-time, time-homogeneous Markov chain where the set of possible states is *V* and the transition probabilities are *T_ij_* = *f_ij_/ _j_′ f_ij_′* . Starting from any initial state *s*_0_, the particle will transit a random sequence of states *s*_0_*, s*_1_*, s*_2_*, …* Because we have assumed the flux network is strongly connected, that sequence will include all states infinitely many times. Therefore, every time the particle visits a state *i*, it must eventually return to state *i*. The sequence of states in between, *s_t_, …, s_t_′* constitutes a cycle. That need not be a simple cycle–it may comprise multiple nested or interleaved simple cycles. However, it can always be reduced to a unique simple cycle by repeatedly removing the earliest simple cycle found within the sequence—we call this the “first closure reduction rule.” For example, a Markov chain state sequence in Figure 1b may be 1*→*4*→*3*→*2*→*3*→*4*→*3*→*1. Metabolite/state 3 is the first to occur twice in the sequence, so we replace the subsequence 3*→*2*→*3 with just a single 3, resulting in the reduced sequence: 1*→*4*→*3*→*4*→*3*→*1. Now 4 is the first to occur twice, so we replace 4*→*3*→*4 with a single 4, resulting in: 1*→*4*→*3*→*1. This is the unique simple cycle to which our original sequence reduces under the first closure reduction rule, and corresponds to EFM 6. In similar fashion, every revisit to any state can be said to have occurred via a particular simple cycle, which of course corresponds to an EFM. Therefore, we propose finding unique EFM weights by setting them proportional to the (steady state) probabilities of our Markovian particle transiting the corresponding simple cycles.

#### 2.2.2 Cycle-history Markov chain

Equating EFM weights with cycle probabilities in the Markov chain in the previous section creates the problem of computing those cycle probabilities which has not been previously addressed in the Markov chain literature. We take the first step in that direction by defining a cycle-history Markov chain (CHMC) and show how to use it to compute the desired probabilities. The CHMC is related to the single particle Markov chain, but its notion of state is expanded to include enough of the history of the process to determine each time the particle finishes transiting a simple cycle. The CHMC is constructed as follows. We first choose an arbitrary metabolite *s*_0_ in the flux network. (Because our interest is in steady state properties of the Markov chain, the choice of *s*_0_ is not important to the final result, although it may result in some differences in the size of the CHMC being constructed). Each state *S* in the CHMC corresponds to an entire simple path *s*_0_, *s*_1_, …, *s_m_*of the Markov chain envisaged in the single particle Markov chain. If there is a reaction transforming metabolite *s_m_* to *s_m_*_+1_, then there is a corresponding transition in the CHMC, but it takes one of two forms. If *s_m_*_+1_ is not among *s*_0_, *s*_1_, … , *s_m_*, then it is simply a transition to the longer simple path–i.e. from the state *S ≡ s*_0_*, s*_1_*, … , s_m_* to a state *S^′^ ≡ s*_0_*, s*_1_*, … , s_m_, s_m_*_+1_. However, if *s_m_*_+1_ is among *s*_0_, *s*_1_, … , *s_m_*, say *s_m_*_+1_ = *s_i_* for 0 *≤ i < m*, then this corresponds to closing a simple cycle, and there is a transition from *S ≡ s*_0_*, s*_1_*, … , s_m_* back to the shorter simple path *S^′^ ≡ s*_0_*, s*_1_*, … , s_i_*. In either case, the probability of that transition in the CHMC is the same as the transition in the Markov chain: *T_S,S_′* = *T_s__m,sm_*_+1_

To make that abstract discussion more concrete, let us first consider the flux network from Figure 1a. Suppose the CHMC for that flux network is initialized at metabolite 1. The chain is “grown” recursively by adding children metabolites 2 and 3, and so forth, until there are four simple paths ending at the special external environment node. The resulting CHMC is shown in Figure 2a with four additional upstream edges transitioning back to metabolite 1 that mark the completion of a simple cycle for each EFM starting and ending at the root node.

**Figure 2:**
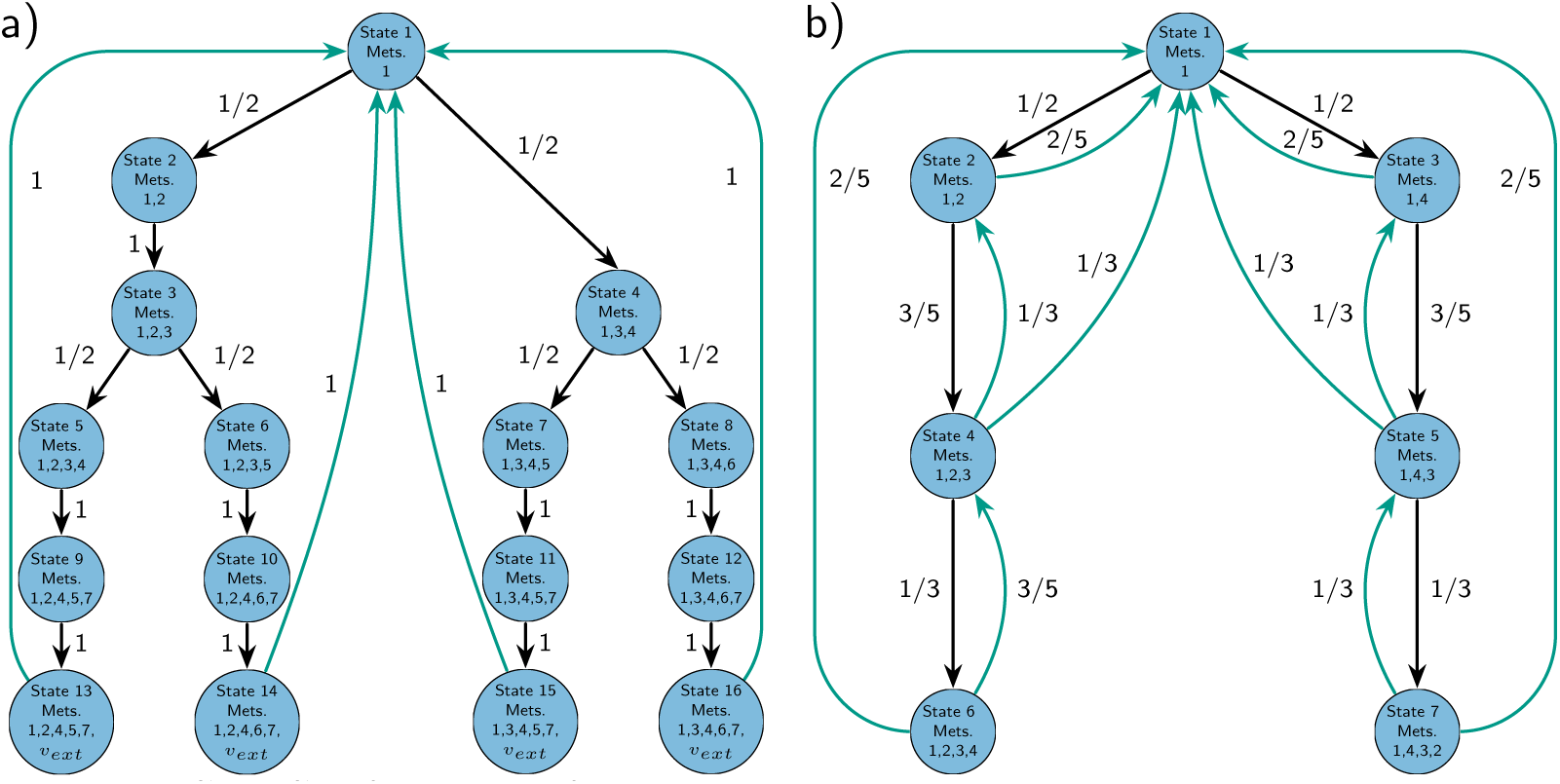
CHMCs of networks from Figure 1 that are arbitrarily rooted on metabolite 1. Nodes and edges denote metabolite sequences and CHMC transition probabilities, respectively. Coloured arrows represent the completion of a simple cycle corresponding to an EFM.

Figure 2b shows the CHMC for the second example network. Every EFM appears as a loop somewhere in the transformed network. EFMs involving the initial metabolite occur as precisely one path from root to leaf and then back up to root, such as EFMs 7 and 8. EFMs not involving the root metabolite also occur somewhere in the CHMC, such as EFM 1 which cycles between metabolites 2 and 1. However, other EFMs may occur in multiple parts of the tree. For instance, EFM 2, which is a loop between metabolites 2 and 3, is represented by two cycles in the CHMC: from state 2 to 4 and back to 2, and also from state 5 to 7 and back to 5.

#### 2.2.3 Computing steady state EFM probabilities

As we have assumed the flux network is strongly connected, the corresponding CHMC will also be strongly connected. This implies the existence of a unique steady state distribution *π* which is the long-term probability of the particle occupying each CHMC state. To compute the steady state probability of EFM

*k*, we identify all transitions in the CHMC that correspond to closing that cycle, *E_k_*. For example, returning to Figure 2b and EFM 2, we have *E*_2_ = *{*(4, 2), (7, 5)*}*, as those two edges close cycles involving the 2-3-2 metabolic loop. For other EFMs, *E_k_* may contain just a single transition. We define the steady state probability of transiting EFM *k* as the sum of the steady state probabilities of those transitions occurring, or:

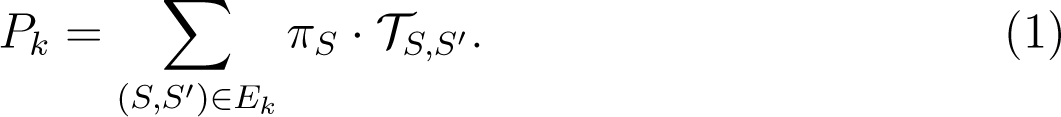

In our proposal, the steady state EFM probabilities *P_k_* are proportional to the desired EFM weights *w_k_* and must be scaled appropriately to account for the absolute magnitude of the fluxes. If *f_total_* = *_i,j_ f_i,j_* is the total flux in the network summed across reactions, and *f_efm_* = ^∑^*_k_ w_k_L_k_* is the total flux obtained by summing EFM weights *w_k_* and their lengths *L_k_*, then our proportionality constant *α*, where *w_k_* = *αP_k_*, should satisfy *f_total_* = *f_efm_* = *α _k_ P_k_L_k_*, so *α* = *f_total_/ _k_ P_k_L_k_*. This scaling ensures not only that the total flux is correct, but that every individual flux in the network is correctly accounted for.

### 2.3 Comparison to optimization-based methods

We benchmarked the Markov model against five optimization-based methods reported in literature. The LP, QP, and MILP objective functions are described in Table 1. Briefly, these approaches optimize either the zero-, one- or two-norm of the EFM weights, possibly multiplied by the EFM lengths, and subject to constraints that ensure the fluxes are correctly reconstructed. EFM enumeration and optimization programs were implemented in Julia (version 1.6.5) using the commercially-licensed Gurobi solver (version 9.1.2) and open-source solvers supported in the Julia JuMP package. In total, EFMs weights for the optimization-based methods were computed using eight LP, two QP, and one MILP solver (see Supplementary information section 2).

**Table 1:** Benchmarked optimization-based approaches

| Description and code | Objective function | Ref. |
| --- | --- | --- |
| Minimize L2 norm (L2) | $\min \sum w_k^2$ | [31] |
| Maximize shortest pathway activity (qSPA) | $\max \sum (L_k w_k)^2$ | [47] |
| Maximize shortest pathway activity (lSPA) | $\max \sum (L_k w_k)$ | [48] |
| Minimize shortest pathway activity (lSPA) | $\min \sum (L_k w_k)$ | [49] |
| Minimize active pathways (milAP) | $\min \sum \beta_k = \begin{cases} 1, & \text{if } w_k > 0 \\ 0 & \text{otherwise.} \end{cases}$ | [32] |

## 3 Results

### 3.1 Markov model uniquely estimates EFM weights in example networks

We first compared the Markov and optimization-based methods on the example networks in Figure 1. In the first example network, the steady state probabilities of the particle occupying external metabolite states *v_ext_*, denoted by the CHMC nodes 13-16 in Figure 2a, were 4.545% because the path probabilities along each prefix were identical. As the probability of completing each simple cycle was 100% from any of those states, the steady state EFM probabilities were 4.545%. Scaling the EFM probabilities to weights yielded the unique solution *w* = [1111]. In contrast, the solutions for the five optimization-based methods, presented in Table 2, returned three feasible EFM weight solutions across all eleven solvers. Of the five objective functions used, minimizing the two QP objective functions, across all solvers, returned only the Markovian solution. However, the LP-based programs returned three solutions across all eight solvers, including the Markovian weights. Given all EFMs were of the same length, neither LP objective function constrained the solution space and could have returned an infinite set of decimal-valued EFM weights. The MILP method was the only one that failed to return the Markovian solution and returned one of two feasible solutions that minimized the number of active EFMs.

**Table 2:** EFM weights from Figure 1a) with solvers shown below.

| EFM | Method |  |  |  |  |  |  |  |  |  |
| --- | --- | --- | --- | --- | --- | --- | --- | --- | --- | --- |
|  | Markov | Min L2 | Max qSPA | Max lSPA |  |  | Min lSPA |  |  | Min milAP |
| 1 | 1 | 1<sup>a</sup> | 1<sup>b</sup> | 2<sup>c</sup> | 0<sup>d</sup> | 1<sup>e</sup> | 2<sup>f</sup> | 0<sup>g</sup> | 1<sup>h</sup> | 2<sup>i</sup> |
| 2 | 1 | 1 | 1 | 0 | 2 | 1 | 0 | 2 | 1 | 0 |
| 3 | 1 | 1 | 1 | 2 | 0 | 1 | 2 | 0 | 1 | 2 |
| 4 | 1 | 1 | 1 | 0 | 2 | 1 | 0 | 2 | 1 | 0 |

<sup>a</sup>COSMO, OSQP; <sup>b</sup>COSMO, OSQP; <sup>c</sup>CDDLib; <sup>d</sup>Gurobi, SCIP, GLPK; <sup>e</sup>OSQP, ECOS, ProxSDP, Tulip; <sup>f</sup>CDDLib;<sup>g</sup>Gurobi, SCIP, GLPK; <sup>h</sup>OSQP, ECOS, ProxSDP, Tulip; <sup>i</sup>Gurobi.

The EFM probabilities were next computed for the second example network shown in Figure 1b. The EFM probabilities for Figure 2b were *P_k_* = [0.17, 0.17, 0.17, 0.17, 0.11, 0.11, 0.043, 0.043] which, when rescaled by proportionality constant *α*, yielded *w_k_* = [1.6, 1.6, 1.6, 1.6, 1.0, 1.0, 0.4, 0.4]. This Markovian solution differed greatly from all five optimization-based methods presented in Table 3. Of the seven feasible solutions returned across all objective functions, none were the Markovian solution and methods that returned only a single solution were different from each other. Collectively, these results demonstrated that the Markovian constraint identified EFM weights that were unique and distinct from the other optimization-based methods, which returned multiple feasible solutions depending on the objective function and solver.

**Table 3:**
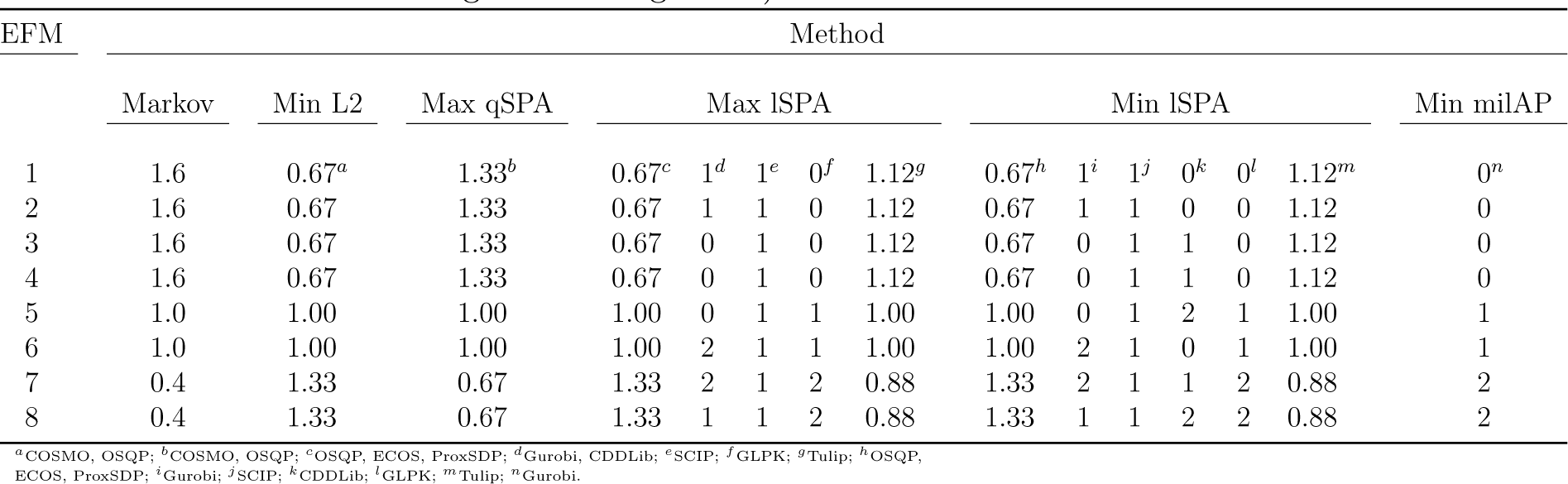
EFM weights from Figure 1b) with solvers shown below.

### 3.2 Application to a sphingolipid network

We next compared the Markov model and optimization-based methods to assign EFM weights in an empirically-validated network of sphingolipid metabolism of non-specific human tissue across nine subcellular compartments [12]. Figure 3 shows the model which consists of 69 reactions across 39 lipid species and kinetic parameters for the wildtype and Alzheimer’s disease condition.

The within-compartment reactions were modelled by Michaelis-Menten kinetics, while inter-compartment transport reactions were modelled by mass-action kinetics. This sphingolipid network is an ideal unimolecular model because reactions either move a lipid family from one compartment to another or transform one lipid family into another by the addition or removal of an N-acyl fatty acid or phosphocholine head group. Although Figure 3 suggests an open-loop network, source reactions 64-68 and sink reaction 6 contained zero-valued kinetic parameters in both conditions and therefore no flux into or out of the system. The kinetic parameter for source reaction 1 was zero in the wildtype condition and set to zero (from *V_m_* = 0.04) in the disease condition to ensure a closed-loop network. Altogether, there were 62 remaining reactions involving 37 lipid species.

**Figure 3:**
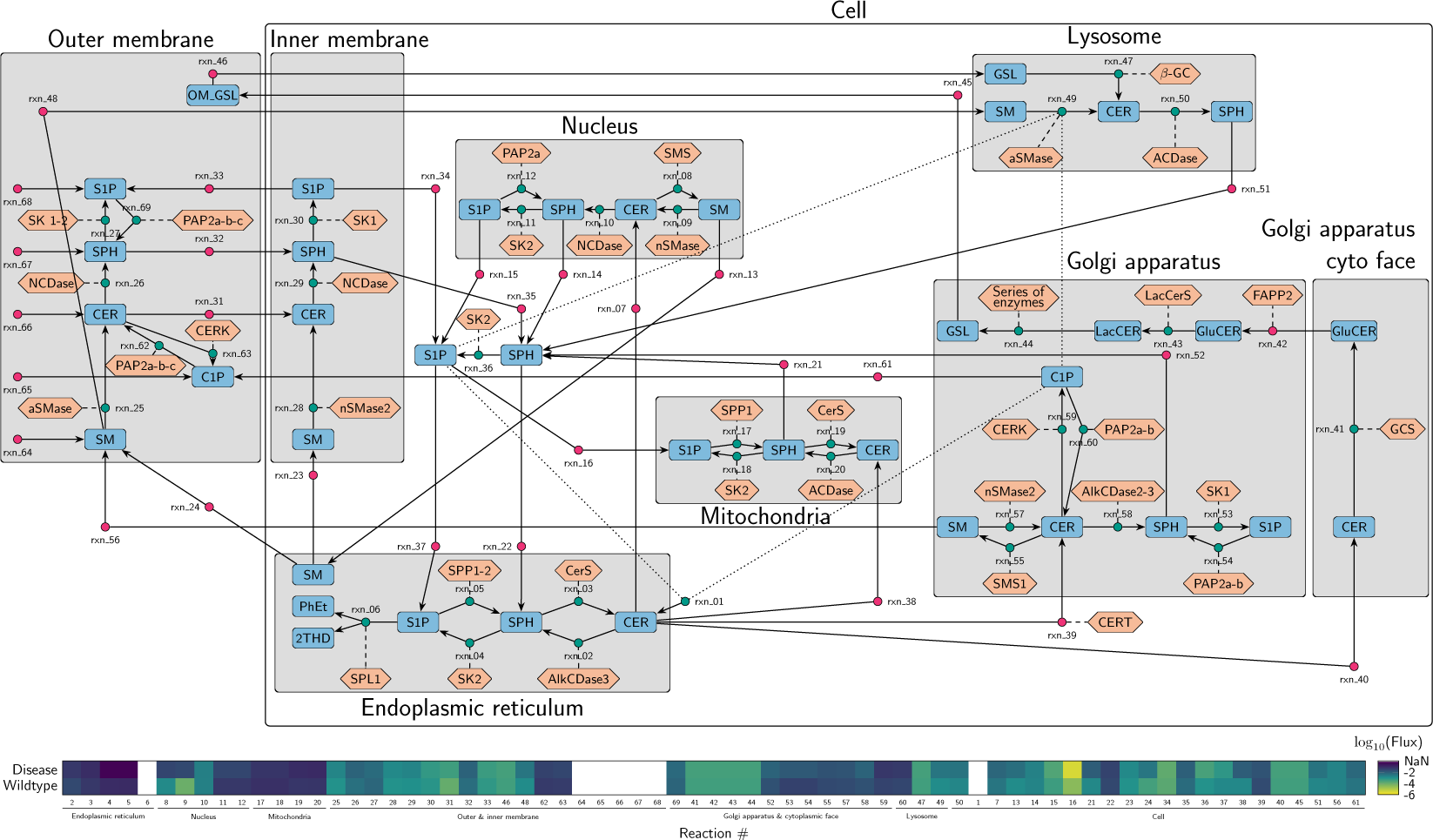
Sphingolipid kinetic model adapted from Wronowska et al. Oval boxes denote lipid families and hexagonal boxes denote enzymes. Teal circles denote transport reactions modelled by mass-action kinetics. Magenta circles denote metabolic reactions modelled by Michaelis-Menten kinetics. Solid lines connect metabolites and reactions, dashed lines denote the reaction enzyme, and dotted lines denote non-competitive inhibition.

The full ordinary differential equation (ODE) models provided in the supplementary material by Wronowska et al. were reproduced with minor corrections and solved to identify the steady state lipid concentrations in both conditions (see Supplementary information section 3). Figure 3 shows the steady state fluxes computed from the kinetic parameters and steady state lipid concentrations of both models.

We next enumerated the EFMs in the sphingolipid network. Of the 55 network EFMs, 11 of them were short, two-step reaction cycles such as ceramide to sphingosine and back to ceramide in the endoplasmic reticulum. Others were longer, including multiple lipid transformations and/or transport between different subcellular locations. The three longest EFMs were 14 steps long and spanned 6 subcellular compartments, although not necessarily the same ones (see Supplementary information section 4). We used our Markovian method as well as the optimization methods described in Table 1 to calculate EFM weights for the healthy and Alzheimer’s disease fluxes.

Figure 4 shows the EFM weights across all tested methods for the wildtype and disease conditions, where all methods, including ours, returned quite different solutions. The greatest agreement in EFM weights, across methods and solvers, was observed for the top 9 EFM weights in the wildtype condition of Figure 4a. These EFMs explained 85.8% of the total network fluxes and consisted nearly exclusively of two-step reversible reactions between lipid families within a compartment. Their weights were consistent across methods because the large fluxes between pairs of reversible reactions can only be explained by these two-step pathways. Beyond these reversible reactions, there was less agreement for the remaining EFMs consisting of longer reactions which varied by up to three orders of magnitude between solvers and across objective functions. As an increase in one EFM weight must be balanced by a decrease in another to satisfy flux conservation, EFMs assigned very large values must force others towards zero. Across all optimization-based methods and solvers, we observed a 32% probability of assigning a zero-valued weight to each of the 55 EFMs. Different objective functions displayed varying tendencies towards sparse solutions with the MILP and maximizing shortest pathway activity by QP functions assigning the most zero-valued weights on average (29 and 26.5). These observations were consistent in the Alzheimer’s disease condition in Figure 4b with high agreement for the same top 9 EFM weights shared in the wildtype condition. These EFMs explained 86.7% of the total network flux with greater disagreement within solvers and across methods for the remaining EFMs. Minimizing the L2 norm, for instance, yielded weights that were up to four magnitudes smaller than the Markov solution or other objective functions. This variability corresponded to a 35% probability of any optimization-based method/solver assigning a zero-valued weight. In contrast, our Markov solution is not based on sparsity or parsimony, so all EFMs receive some non-zero weight, regardless of their length or how much flux they explain in the network. We observed that the largest 13 EFM weights explained 95% of the fluxes in both conditions and that 11 of them were shared between wildtype and Alzheimer’s disease (see Supplementary information section 5). One of the two largest EFMs unique to the wildtype and Alzheimer’s disease were two-step reactions cycling sphingosine and sphingosine-1-phosphate in the outer membrane and sphingomyelin to ceramide in the nucleus, respectively.

**Figure 4:**
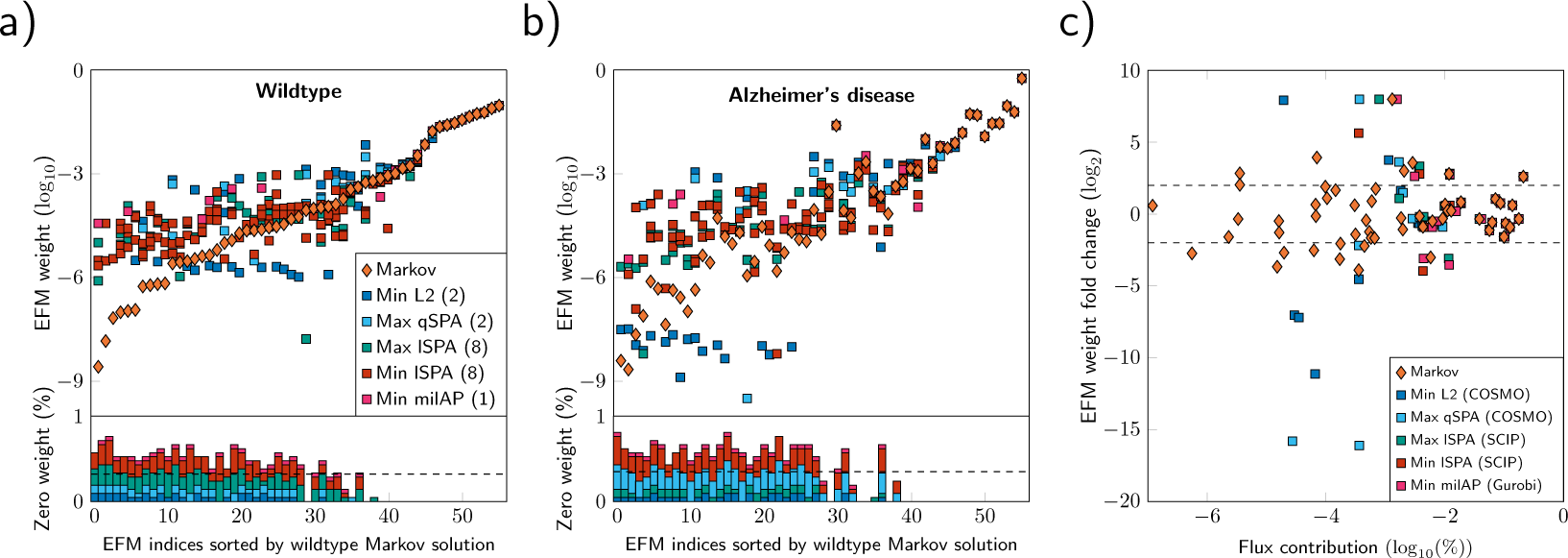
EFM weights under the Markov constraint versus five objective functions from optimization-based methods across all solvers for the a) wildtype, b) disease condition, c) fold change with respect to wildtype with a representative solver from each method The optimization-based method codes are described in Table 1. Numbers in parentheses correspond to the solvers used for each mathematical optimization. Histogram bars show the frequency of zero-valued EFM weights from the optimization-based methods normalized across all 21 objective functions and solvers.

Given the majority of network fluxes across both conditions were explained by the same 11 EFMs, we next investigated whether metabolic differences were explainable through other differentially active EFMs. Figure 4c shows the fold change between the Alzheimer’s disease and wildtype condition EFM weights as a function of the total explained flux in the wildtype condition. Weights were selected from the objective function solver that best reconstructed the individual and total fluxes across both networks (see Supplementary information sections 6 & 7). The top 9 EFM weights explaining the majority of the network fluxes are located on the right-most cluster of Figure 4c where there is high agreement across all methods. Only one of those EFMs showed over four-fold increase in the Alzheimer’s disease versus wildtype condition. This two-step EFM pathway corresponded to the remodelling of sphingosine and sphingosine-1-phosphate in the endoplasmic reticulum. According to the Markov solution, we observed extreme fold changes (*|*log_2_ *FC| ≥* 4) for 17 EFMs that explained between 10*^−^*^6^ and 0.1% of the total network flux. In contrast, there were fewer fold changes for the optimization-based methods due to the sparsity of EFM weights for the disease and wildtype conditions. Between 25 and 34 undefined fold changes were observed across all five optimization methods because these methods assigned a zero-valued EFM weight to one or both conditions. Furthermore, the optimization methods also showed extreme fold changes for EFMs that explained far less flux than our Markov solution. Of the 6 to 10 extreme fold changes observed across all LP/MILP/QP methods, their EFM weights only explained between 0.0001% to 0.001% of the total wildtype fluxes.

## 4 Conclusion

In this article, we explored the computational problem of identifying EFM weights explaining steady state fluxes in a metabolic network. Although this problem has mainly been addressed by optimization-based methods, we showed how different objective functions may nonetheless fail to properly constrain the solution space of EFM weights. We then proposed a Markovian constraint, in which EFM weights are made proportional to the probability of a metabolite “particle” travelling along different paths through the network. Our Markovian approach results from a natural, biophysical assumption, which may offer greater accuracy to real metabolic networks.

We applied our Markov method to identify differentially active EFM weights in a model of sphingolipid metabolism in Alzheimer’s disease and compared the results against optimization-based methods. While all methods strongly agreed on the top 9 (16%) EFM weights, the Markov and objective functions across all solvers greatly differed for the remaining EFMs. Upon inspection, we found that EFMs with the largest weights and consensus across flux decomposition methods corresponded to two-step remodelling cycles between pairs of lipid families, while EFMs with smaller weights typically involved more reactions and/or spanned compartments.

While we developed our CHMC method to uniquely estimate EFM weights from metabolic flux networks, we highlight algorithmic applications for other network flow problems in biology. Ion channel behaviour or protein folding dynamics may be characterized by the cyclic flow of conformational changes of the atom positions and momenta [50, 51], and these simple cycle probabilities may be uniquely computed by our CHMC method. Flow decomposition ambiguity also occurs in RNA quantification algorithms where the same set of reads mapped to a splice graph may be explained by more than one set of transcript isoforms, even under parsimonious constraints [52, 53].

There exist several avenues of future work to extend our Markov chain model to identify EFM weights. First, our method is only applicable to metabolic networks of unimolecular reactions and our methodology must be expanded to account for networks containing higher order reactions. Second, algorithmic developments are required to scale our Markov method to genomewide networks containing millions of EFMs. While the Julia code we provide can compute two to three thousand EFM weights in seconds, enumerating the CHMC state space is prohibitively memory intensive in larger-scale networks. Third, we anticipate our work will encourage differential analysis of EFM weights to quantify metabolic changes across conditions. Previous analyses have typically computed EFM weights using a variety of objective functions and used a consensus-based approach to identify active EFMs [31, 47]. As our method uniquely identifies EFM weights, we foresee a deeper analysis of EFM weights and their incorporation in statistical learning models to study differential metabolism.

## 5 Data availability

Our CHMC implementation is publicly available at https://github.com/jchitpin/MarkovWeightedEFMs.jl. The codes and datasets to reproduce all figures are available at https://github.com/jchitpin/reproduce-efm-paper-2023.

## 6 Supplementary information

Supplementary information are included in this manuscript submission.

## 7 Competing interests

No competing interest is declared.

## 8 Author contributions statement

J.G.C. and T.J.P conceived the study, J.G.C. conducted the experiments and analyzed the results. J.G.C. and T.J.P. wrote and reviewed the manuscript.

## Acknowledgments

We acknowledge the support of the Natural Sciences and Engineering Research Council of Canada (NSERC), Discovery grant RGPIN-2019-0660 to T.J.P. J.G.C. was supported by an NSERC CREATE Matrix Metabolomics Scholarship and an NSERC Alexander Graham Bell Canada Graduate Scholarship.

## Notes

### Competing Interest Statement

The authors have declared no competing interest.

### Summary of Updates

Revision of presentation, mathematical correctness proof, and more detailed analysis of Alzheimer's model.

