## supplementary for "A Markov constraint to uniquely identify elementary flux mode weights in unimolecular metabolic networks"

**2 List of optimization-based method solvers** **8**

**3 Sphingolipid kinetic model (reproduced with minor corrections)** **9**

**4 Sphingolipid network elementary flux modes** **13**

**5 Markov weights for the wildtype and Alzheimer's disease sphingolipid network** **14**

**6 Individual flux reconstruction error across methods** **15**

**7 Total flux reconstruction error across methods** **15**

### 1 Proof of correctness of the CHMC algorithm

In this section, we establish the correctness of the CHMC algorithm described in the main text. By correctness, we mean that it calculates EFM weights that attribute the correct amount of flux for every reaction in the network. The first section below reprises some definitions from the main text. The next section establishes some lemmas about the steady state properties of the Markov chain that tracks a single particle moving around the network. We then establish some lemmas about steady state properties of the CHMC. And in the final section, we complete the proof of the theorem by showing that the fluxes are correctly explained.

#### 1.1 Flux network basics

A *flux network* is a triple $G = (V, E, f)$, where $V = \{1, 2, \ldots, n\}$ is a set of nodes, $E \subseteq V \times V$ is a set of directed edges, and $f$ are positive edge weights. We assume no node has an edge directly back to itself: $(i, i) \notin E$ for all $i \in V$. $f$ may be defined either as a function $f : E \to \Re^+$, or as a function $f : V \times V \to \Re^{\geq 0}$ where $f_{i,j} > 0 \iff (i, j) \in E$.

We restrict attention to strongly-connected flux networks, meaning that every node is reachable from every other. We also restrict attention to steady-state flux networks, meaning that the flux into any node equals the flux out of the node: for all $i \in V$, $\sum_{j \neq i} f_{i,j} = \sum_{j \neq i} f_{j,i}$. We call this the flux *at* node $i$, and overload the $f$ notation to denote the flux at node $i$ as $f_i$.

We define the *total network flux* as $f_{tot} = \sum_{i,j} f_{i,j}$, and we define the *normalized fluxes* as $f^n_{i,j} = f_{i,j}/f_{tot}$. Obviously, if a set of elementary flux mode (EFM) weights explains all the normalized fluxes $f^n$ in a network, then $f_{tot}$ times those weights explains all unnormalized fluxes $f$.

Finally, as already implied by the definitions above, we restrict attention to closed flux networks, meaning there is no flux into or out of the system. However, this is a matter merely of mathematical convenience. Any open flux network, where sources nodes produce flux and sink nodes consume flux, can be trivially handled by our approach as well. One simply represents the "exterior" of the network by a special node, and connects flux providing and consuming edges to the source and sink nodes respectively.

In closed flux networks, the EFMs are the simple cycles of in network–paths that begin and end at the same node, with no nodes repeated in between. We use $C = \{c_1, c_2, \ldots, c_k\}$ to denote this set of cycles. Any particular cycle is of the form $(v_1, v_2, \ldots, v_{m-1}, v_m)$, where $v_1 = v_m \in V$, but no other $v_i$ are equal to each other.

#### 1.2 Markov chain steady state properties

Based on the flux network $G$ we define a Markov chain $M = (S, T, s_0)$ where the state space is the same as the nodes of the flux network: $S = V$. The state-to-state transition probabilities $T$ are proportional to the fluxes in the following sense: where $s_t$ denotes a random variable representing the state of the chain at time $t$, $T_{i,j} \equiv Pr(s_{t+1} = j | s_t = i) = f_{i,j} / \sum_{j'} f_{i,j'}$. We arbitrarily choose the initial state of the Markov chain to be $s_0 = 1$.

Because the chain network $G$ is strongly connected, the Markov chain $M$ is irreducible. It therefore has a unique steady state distribution $\pi$ satisfying $\pi \cdot T = \pi$. Indeed, the steady state probabilities are just the normalized fluxes at the nodes:

**Lemma 1** *The steady state distribution $\pi$ of the Markov chain $M$ is equal to the vector of normalized state fluxes $f^n$ of the flux network $G$: $\pi = f^n$.*

Proof: Let $f^n$ be a row vector of the fluxes at the states $1, \ldots, n$, and consider the $j^{th}$ column of the Markov transition matrix, which we will denote $T_{\cdot,j}$. Then:

$$f^n \cdot T_{\cdot,j} = \sum_i f_i^n \cdot T_{i,j} = \sum_i f_i^n \cdot \frac{f_{i,j}}{f_i^n} = \sum_i f_{i,j}^n = f_j^n$$

Since this is true for any $j$, then we have $f^n \cdot T = f^n$. Note also that $\sum_i f^n = \sum_{i,j} f_{i,j}^n = \sum_{i,j} f_{i,j}/f_{tot} = f_{tot}/f_{tot} = 1$. Since $\pi$ is the unique steady state vector satisfying $\pi \cdot T = \pi$ and $\sum_i \pi = 1$, and since $f^n$ satisfies the same two properties, then $\pi = f^n$.

We define the probabilistic flux $\pi_{i,j}$ of the Markov chain as the steady state frequency with which transitions from state $i$ to state $j$ occur, or equivalently, $\pi_{i,j} = \pi_i \cdot T_{i,j}$. It follows immediately that $\pi_{i,j} = f_{i,j}^n$, which we state in the lemma below without further argument.

**Lemma 2** *The probabilistic fluxes $\pi_{i,j}$ of the Markov chain $M$ are equal to the normalized fluxes $f_{i,j}^n$ of the flux network $G$.*

##### 1.3 Cycle-History Markov chain steady state properties

The cycle-history Markov chain (CHMC) is a Markov chain $M^c = (S^c, T^c, s_0^c)$ that is constructed based on the Markov chain $M$. Every state $s \in S^c$ in the CHMC corresponds to a simple path of chain $M$ starting from its initial state $s_0$. That is, $s \equiv (s_0, s_1, \ldots, s_m)$ for some $m \geq 0$, where all $s_i \in S$, $s_0 = 1$, and $s_i \neq s_j$ for all $0 \leq i < j \leq m$. The state set $S^c$ corresponds to all possible simple paths in $M$, where "possible paths" means paths with strictly positive probability. The initial state of the CHMC corresponds to the trivial zero-step path starting at $M$'s initial state: $s_0^c \equiv (s_0)$. For every CHMC state $s \equiv (s_0, s_1, \ldots, s_m)$ and for every Markov chain state $j$ that can be transitioned to from $s_m$ (i.e. every $j$ where $T_{s_m,j} > 0$), there is a corresponding transition in the CHMC with the same probability. If state $j$ is not on the path $(s_0, s_1, \ldots, s_m)$, then the transition is to a CHMC state $s' \equiv (s_0, s_1, \ldots, s_m, j)$. If state $j$ is on the path $(s_0, s_1, \ldots, s_m)$, say at $s_k = j$, then it is a transition to the CHMC state corresponding to the shorter path $s' \equiv (s_0, s_1, \ldots, s_k)$. This latter type of transition is called a cycle-/loop-/EFM-closing transition, because it corresponds to the Markov chain finishing transiting a cycle—specifically, the cycle given by the latter portion of $s$'s path: $(s_k, s_{k+1}, \ldots, s_m, s_k)$. In the CHMC, it is self-evident that every state is reachable from every other state. Therefore, the CHMC has a unique steady state distribution $\pi^c$ satisfying $\pi^c \cdot T^c = \pi^c$.

Whereas the CHMC construction "blows up" the Markov chain $M$'s state space into a much larger state space of all possible paths, it is useful to consider a sort of reverse transformation where we aggregate together different states of the CHMC based on the final state of their path. That is, define a function $A : S^c \mapsto S$ where for $s \equiv (s_0, s_1, \ldots, s_m)$, we have $A(s) = s_m$. Recalling that the state set of the Markov chain $M$, and the original flux network $G$, are just the numbers $1, 2, \ldots, n$, then $A$ is just a mapping from $S^c$ to the numbers 1 to $n$. For convenience, let us also define the set of CHMC states mapping to each Markov state $i$: $S^c_i = \{s \in S^c : A(s) = i\}$.

We define the aggregated CHMC steady state distribution as $\pi^A$ via: $\pi^A_i = \sum_{s \in S^c_i} \pi^c_s$. In words, $\pi^A$ is a vector where the $i^{th}$ element is the sum of steady state probabilities of all CHMC state corresponding to paths ending at $i$. Our first main observation about the CHMC is that the aggregated steady state probabilities are equal to the Markov chain steady states.

**Lemma 3** *Where $\pi$ is the steady state distribution of the Markov chain $M$, and where $\pi^A$ are the aggregated steady state probabilities of the CHMC $M^c$: $\pi^A = \pi$.*

This makes intuitive sense. The CHMC is basically keeping multiple "copies" of a Markov state $i$ based on the path followed to reach state $i$. However, $i$ always has to be reached by some path. Therefore, by summing over all of those paths, we should arrive at the total probability of being in state $i$. However, we can argue more formally as follows.

Proof: Consider any Markov state $i$ and consider any CHMC state corresponding to a path ending at that state, $s \in S^c_i$. Let $T^c_{\cdot,s}$ denote the column of the CHMC transition matrix corresponding to transitions into state $s$. Then we have

$$\pi^c_s = \pi^c \cdot T^c_{\cdot,s} = \sum_{s' \in S^c} \pi^c_{s'} \cdot T^c_{s',s}$$

The summation over $s'$ can alternatively be written as a double sum over Markov chain states $j$ and CHMC states corresponding to paths ending at $j$:

$$\pi^c_s = \sum_{j \in S} \sum_{s' \in S^c_j} \pi^c_{s'} \cdot T^c_{s',s}$$

We can further sum both sides of the equation over all CHMC states $s''$ in the same partition as $s$:

$$\sum_{s'' \in S^c_i} \pi^c_{s''} = \sum_{s'' \in S^c_i} \sum_{j \in S} \sum_{s' \in S^c_j} \pi^c_{s'} \cdot T^c_{s',s''}$$

We recognize the left hand side as the aggregated CHMC probability of states corresponding to paths ending at state $i$. Meanwhile, the right hand side can be rewritten by reordering the sums.

$$\pi^A_i = \sum_{j \in S} \sum_{s' \in S^c_j} \sum_{s'' \in S^c_i} \pi^c_{s'} \cdot T^c_{s',s''}$$

The double summation over $s'$ and $s''$ is not really a double summation, in the sense that for any $s' \in S^c_j$, and assuming a transition to some $s'' \in S^c_i$ is possible at all (i.e. assuming $T_{j,i} > 0$) then there will be one and precisely one $s''$ to which $s'$ can transition. All other transitions from $s'$ to other elements of $S^c_i$ will have probability zero. To be precise, if $s''$ is either a shorter part of the path represented by $s'$, or if $s''$ is a one-step extension of the path represented by $s'$, then we will have $T^c_{s',s''} = T_{j,i}$. Otherwise, we will have $T^c_{s',s''} = 0$. Therefore, we can rewrite as:

$$\pi^A_i = \sum_{j \in S} \sum_{s' \in S^c_j} \pi^c_{s'} \cdot T_{j,i}$$

From this, we deduce that:

$$\pi_i^A = \sum_{j \in S} \pi_j^A \cdot T_{j,i}$$

Because this is true for any Markov state $i$, we must have $\pi^A = \pi^A \cdot T$, meaning that $\pi^A$ is the (unique) steady state distribution for the Markov chain $M$, or in other words $\pi = \pi^A$, establishing the lemma.

A similar result is true for the aggregated probabilistic fluxes of the CHMC. The probabilistic flux from CHMC state $s$ to state $s'$ is $\pi^c_{s,s'} = \pi^c_s \cdot T^c_{s,s'}$. We then define the aggregated probabilistic fluxes for any $i, j \in S$, as $\pi^A_{i,j} = \sum_{s \in S^c_i} \sum_{s' \in S^c_j} \pi^c_{s,s'}$.

**Lemma 4** *The aggregated probabilistic fluxes $\pi^A_{i,j}$ of the CHMC $M^c$ are equal to the probabilistic fluxes $\pi_{i,j}$ of the Markov chain $M$, which are equal to the normalized fluxes $f^n_{i,j}$ of the flux network $G$.*

Proof: We start from the definition of the aggregated probabilistic flux:

$$\pi^A_{i,j} = \sum_{s \in S^c_i} \sum_{s' \in S^c_j} \pi^c_{s,s'} = \sum_{s \in S^c_i} \sum_{s' \in S^c_j} \pi^c_s \cdot T^c_{s,s'} = \sum_{s \in S^c_i} \pi^c_s \cdot \sum_{s' \in S^c_j} T^c_{s,s'}$$

As explained in the previous section, the apparent double sum over $s$ and $s'$ really only has one non-zero term per $s$, because there is only one (at most) non-zero transition from any element of $S^c_i$ to any element of $S^c_j$. Therefore, we can write:

$$\pi^A_{i,j} = \sum_{s \in S^c_i} \pi^c_s \cdot T_{i,j} = \pi^A_i \cdot T_{i,j} = \pi_i \cdot T_{i,j} = \pi_{i,j}$$

which completes the lemma.

##### 1.4 Explaining fluxes

We begin by establishing a means by which CHMC probabilistic fluxes $\pi^c_{s,s'}$ (not the aggregated ones) can be "explained" in terms of weighted cycles, and then we extend that result to show that a simple aggregation and rescaling of those same weights explains the fluxes $f_{i,j}$ in the original flux network $G$.

We first show that the CHMC probabilistic fluxes can be explained by interpreting them as a flux network. Specifically, consider the flux network $G' = (V', E', f')$ where the nodes are the same as the CHMC states: $V' = S^c$. The transitions are $(s, s') \in E'$ iff $T^c_{s,s'} > 0$. And the fluxes are $f' = \pi^c$. Because this is a flux network, we know that the fluxes can be explained by a weighted combination of elementary flux modes. What are the EFMs of $G'$? First, $G'$ is a closed network, and so the EFMs are a set of cycles, $C' = \{c'_1, c'_2, \ldots, c'_{k'}\}$. However, it is a very special set of cycles. By construction the CHMC transitions form a tree, with the exception of the loop-closing transition. Therefore, every cycle $c'$ contains precisely one loop-closure edge. And conversely, no other cycle $c''$ includes that same loop-closure edge. Therefore, let us denote those loop closure edges as $(s_i, s'_i) \in E'$ for $1 \leq i \leq k'$, one for each cycle. When an EFM is the one and only EFM to pass through a certain edge in a flux network, then the flux (or weight) of that EFM must equal that flux of that edge—because there is no other way to explain the flux on that edge. And since every EFM has one such edge, we know therefore that there can only be one weighting of the EFMs that explains the probabilistic flux $f'$, namely: EFM $i$ must be given weight $f_{s_i,s'_i} = \pi^c_{s_i,s'_i}$. Therefore, we have the following lemma.

**Lemma 5** *There is a set of non-negative weights $w'_1, w'_2, \ldots, w'_{k'}$ that fully explain (probabilistic) fluxes $f'$ in the sense that: For all $s, s' \in V'$:*

$$f'_{s,s'} = \sum_{i=1}^{k'} w'_i \cdot \begin{cases} 1 & \text{if } (s,s') \in c'_i \\ 0 & \text{otherwise.} \end{cases}$$

*In particular, the weights are $w'_i = \pi^c_{s_i,s'_i}$ where $(s_i, s'_i)$ is the loop-closure edge in the $i^{th}$ cycle (a.k.a. EFM) $c'_i$.*

Finally, we connect the weights of the previous lemma to the original flux network $G$. As stated in the main manuscript, by construction of the Markov chain $M$, the CHMC $M^c$, and the flux network $G'$, every EFM in $G'$ corresponds to an EFM in $G$. However, as also demonstrated in the main text, an EFM in $G$ can correspond to more than one EFM in $G'$. In essence, $G'$ can have multiple "copies" of the same EFM in $G$, prefaced by different paths reaching that EFM. Therefore, where $C = \{c_1, c_2, \ldots, c_k\}$ are the EFMs of $G$ and $C' = \{c'_1, c'_2, \ldots, c'_{k'}\}$ are the EFMs of $G$, we can define a many-to-one mapping relating the two: $Z : C' \mapsto C$. And let us define the aggregated weights $w^Z_i$ for $1 \leq i \leq k$ as: $w^Z_i = \sum_{\{c'_j : Z(c'_j) = c_i\}} w'_j$, where $w'_j$ are the EFM weights for $G'$ of the previous lemma. And finally, let us define the scaled aggregated weights as $w_1, \ldots, w_k$ where $w_i = f_{tot} \cdot w^Z_i$. Then we arrive at our main theorem.

**Theorem 1** *The weights $w_1, w_2, \ldots, w_k$ for the EFMs $C$ of flux network $G$ fully explain the normalized fluxes $f$ in the sense that: For all $i, j \in V$:*

$$f_{i,j} = \sum_{l=1}^{k} w_l \cdot \begin{cases} 1 & \text{if } (i,j) \in c_l \\ 0 & \text{otherwise.} \end{cases}$$

Proof: Consider any $i, j \in V$. Starting from the right hand side of the equation in the theorem, we have:

$$\begin{aligned} & \sum_{l=1}^{k} w_l \cdot \begin{cases} 1 & \text{if } (i,j) \in c_l \\ 0 & \text{otherwise.} \end{cases} \\ = & \sum_{l=1}^{k} f_{tot} \cdot w^Z_l \cdot \begin{cases} 1 & \text{if } (i,j) \in c_l \\ 0 & \text{otherwise.} \end{cases} \\ = & \sum_{l=1}^{k} f_{tot} \cdot \left( \sum_{c'_j : Z(c'_j) = c_l} w'_j \right) \cdot \begin{cases} 1 & \text{if } (i,j) \in c_l \\ 0 & \text{otherwise.} \end{cases} \end{aligned}$$

The double sum is, equivalently, a sum over all cycles $c'$ of the CHMC $M^c$ (or network $G'$). However, the 0/1 condition should be 1 only for those cycles containing what amounts to an $i$ to $j$ transition in the original Markov chain $M$:

$$\begin{aligned}
&= \sum_{m=1}^{k'} f_{tot} \cdot w'_m \cdot \begin{cases} 1 & \text{if } \exists s \in S_i^c, s' \in S_j^c \ s.t. \ (s, s') \in c'_m \\ 0 & \text{otherwise.} \end{cases} \\
&= f_{tot} \sum_{m=1}^{k'} w'_m \sum_{s \in S_i^c, s' \in S_j^c} \begin{cases} 1 & \text{if } (s, s') \in c'_m \\ 0 & \text{otherwise.} \end{cases} \\
&= f_{tot} \sum_{s \in S_i^c, s' \in S_j^c} \sum_{m=1}^{k'} w'_m \begin{cases} 1 & \text{if } (s, s') \in c'_m \\ 0 & \text{otherwise.} \end{cases} \\
&= f_{tot} \sum_{s \in S_i^c, s' \in S_j^c} f'_{s,s'}
\end{aligned}$$

where the last step follows from Lemma 5. Then:

$$\begin{aligned}
&= f_{tot} \sum_{s \in S_i^c, s' \in S_j^c} \pi^c_{s,s'} \\
&= f_{tot} \cdot \pi^A_{i,j} \\
&= f_{tot} \cdot \pi_{i,j}
\end{aligned}$$

where the last step comes from Lemma 4. Then by Lemma 2 we have:

$$\begin{aligned}
&= f_{tot} \cdot f^n_{i,j} \\
&= f_{tot} \cdot \frac{f_{i,j}}{f_{tot}} \\
&= f_{i,j}
\end{aligned}$$

And so the theorem is proved.

#### 2 List of optimization-based method solvers

Table S1: List of solvers for each optimization-based method. All except for Gurobi are open-source and did not require a manual installation step. See `https://jump.dev/JuMP.jl/stable/installation/#Supported-solvers` for more details.

| | COSMO | OSQP | Gurobi | SCIP | CDDLib | ECOS | GLPK | ProxSDP | Tulip |
|---|---|---|---|---|---|---|---|---|---|
| Minimize L2 | ✓ | ✓ | | | | | | | |
| Maximize qSPA | ✓ | ✓ | | | | | | | |
| Maximize lSPA | | ✓ | ✓ | ✓ | ✓ | ✓ | ✓ | ✓ | ✓ |
| Minimize lSPA | | ✓ | ✓ | ✓ | ✓ | ✓ | ✓ | ✓ | ✓ |
| Minimize milAP | | | ✓ | | | | | | |

### 3 Sphingolipid kinetic model (reproduced with minor corrections)

Table S2: Wildtype and Alzheimer's disease reaction parameters in the sphingolipid kinetic model. Inhibition parameters were not provided by Wronowska *et al.* and were manually estimated to satisfy flux conservation and steady state lipid concentrations in the wildtype model. ($Ki_{S1P} = 0.0356898872566651$ and $Ki_{C1P} = 0.1784494362833260$)

| # | Reaction | Flux equation | Wildtype | | | Alzheimers | | |
|---|---|---|---|---|---|---|---|---|
| | | | Km (nmol/mg) | Vm (nmol/min/mg) | k (1/min) | Km (nmol/mg) | Vm (nmol/min/mg) | r (1/min) |
| 1 | → ER.CER | $\frac{Vm_1}{(1+C.S1P/Ki_{S1P})\times(1+GA.C1P/Ki_{C1P})}$ | – | 0 | – | – | 0 | – |
| 2 | ER.CER → ER.SPH | $\frac{Vm_2\times ER.CER}{Km_2+ER.CER}$ | 0.08100 | 0.48000 | – | 0.08100 | 0.32000 | – |
| 3 | ER.SPH → ER.CER | $\frac{Vm_3\times ER.SPH}{Km_3+ER.SPH}$ | 0.17100 | 2.40000 | – | 0.17100 | 2.40000 | – |
| 4 | ER.SPH → ER.S1P | $\frac{Vm_4\times ER.SPH}{Km_4+ER.SPH}$ | 0.00340 | 0.17500 | – | 0.00340 | 0.87500 | – |
| 5 | ER.S1P → ER.SPH | $\frac{Vm_5\times ER.S1P}{Km_5+ER.S1P}$ | 0.03850 | 3.10000 | – | 0.03850 | 3.10000 | – |
| 6 | ER.S1P → ER.PhET + 2THD | $\frac{Vm_5\times ER.S1P}{Km_5+ER.S1P}$ | 0.03500 | 0 | – | 0.03500 | 0 | – |
| 7 | ER.CER → N.CER | $k_7 \times ER.CER - r_7 \times N.CER$ | – | – | 0.80000 | – | – | 0.80000 |
| 8 | N.CER → N.SM | $\frac{Vm_8\times N.CER}{Km_8+N.CER}$ | 0.15500 | 0.10900 | – | 0.15500 | 0.10900 | – |
| 9 | N.SM → N.CER | $\frac{Vm_9\times N.SM}{Km_9+N.SM}$ | 0.12600 | 0.00113 | – | 0.12600 | 0.10000 | – |
| 10 | N.CER → N.SPH | $\frac{Vm_{10}\times N.CER}{Km_{10}+N.CER}$ | 0.06010 | 0.06800 | – | 0.06010 | 0.00453 | – |
| 11 | N.SPH → N.S1P | $\frac{Vm_{11}\times N.SPH}{Km_{11}+N.SPH}$ | 0.00340 | 0.10000 | – | 0.00340 | 0.05000 | – |
| 12 | N.S1P → N.SPH | $\frac{Vm_{12}\times N.S1P}{Km_{12}+N.S1P}$ | 0.02500 | 1.24000 | – | 0.02500 | 1.24000 | – |
| 13 | N.SM → ER.SM | $k_{13} \times N.SM - r_{13} \times ER.SM$ | – | – | 0.12000 | – | – | 0.12000 |
| 14 | N.SPH → C.SPH | $k_{14} \times N.SPH - r_{14} \times C.SPH$ | – | – | 0.50000 | – | – | 0.50000 |
| 15 | N.S1P → C.S1P | $k_{15} \times N.S1P - r_{15} \times C.S1P$ | – | – | 0.44000 | – | – | 0.44000 |
| 16 | C.S1P → M.S1P | $k_{16} \times C.S1P - r_{16} \times M.S1P$ | – | – | 0.25000 | – | – | 0.25000 |
| 17 | M.S1P → M.SPH | $\frac{Vm_{17}\times M.S1P}{Km_{17}+M.S1P}$ | 0.03850 | 3.00000 | – | 0.03850 | 3.00000 | – |
| 18 | M.SPH → M.S1P | $\frac{Vm_{18}\times M.SPH}{Km_{18}+M.SPH}$ | 0.00340 | 0.15000 | – | 0.00340 | 0.07000 | – |
| 19 | M.SPH → M.CER | $\frac{Vm_{19}\times M.SPH}{Km_{19}+M.SPH}$ | 0.00250 | 0.10000 | – | 0.00250 | 0.10000 | – |
| 20 | M.CER → M.SPH | $\frac{Vm_{20}\times M.CER}{Km_{20}+M.CER}$ | 0.14900 | 2.27000 | – | 0.14900 | 2.00000 | – |
| 21 | M.SPH → C.SPH | $k_{21} \times M.SPH - r_{21} \times C.SPH$ | – | – | 0.43000 | – | – | 0.43000 |
| 22 | C.SPH → ER.SPH | $k_{22} \times C.SPH - r_{22} \times ER.SPH$ | – | – | 23.0000 | – | – | 23.0000 |
| 23 | ER.SM → IM.SM | $k_{23} \times ER.SM - r_{23} \times IM.SM$ | – | – | 0.01000 | – | – | 0.01000 |
| 24 | ER.SM → OM.SM | $k_{24} \times ER.SM - r_{24} \times OM.SM$ | – | – | 0.00600 | – | – | 0.01500 |
| 25 | OM.SM → OM.CER | $\frac{Vm_{25}\times M.CER}{Km_{20}+M.CER}$ | 0.04550 | 0.00100 | – | 0.04550 | 0.00200 | – |
| 26 | OM.CER → OM.SPH | $\frac{Vm_{26}\times OM.CER}{Km_{26}+OM.CER}$ | 0.06010 | 0.12000 | – | 0.06010 | 0.08000 | – |
| 27 | OM.SPH → OM.S1P | $\frac{Vm_{27}\times OM.SPH}{Km_{27}+OM.SPH}$ | 0.03400 | 0.11000 | – | 0.03400 | 0.05500 | – |
| 28 | IM.SM → IM.CER | $\frac{Vm_{28}\times IM.SM}{Km_{28}+IM.SM}$ | 0.14800 | 0.00200 | – | 0.14800 | 0.00250 | – |
| 29 | IM.CER → IM.SPH | $\frac{Vm_{29}\times IM.CER}{Km_{29}+IM.CER}$ | 0.06010 | 0.03000 | – | 0.06010 | 0.02000 | – |
| 30 | IM.SPH → IM.S1P | $\frac{Vm_{30}\times IM.SPH}{Km_{30}+IM.SPH}$ | 0.00506 | 0.00300 | – | 0.00506 | 0.00150 | – |
| 31 | OM.CER → IM.CER | $k_{31} \times OM.CER - r_{31} \times IM.CER$ | – | – | 1.00000 | – | – | 1.00000 |
| 32 | OM.SPH → IM.SPH | $k_{32} \times OM.SPH - r_{32} \times IM.SPH$ | – | – | 1.00000 | – | – | 1.00000 |
| 33 | IM.S1P → OM.S1P | $k_{33} \times IM.S1P - r_{33} \times OM.S1P$ | – | – | 1.00000 | – | – | 1.00000 |
| 34 | IM.S1P → C.S1P | $k_{34} \times IM.S1P - r_{34} \times C.S1P$ | – | – | 0.40000 | – | – | 0.40000 |
| 35 | IM.SPH → C.SPH | $k_{35} \times IM.SPH - r_{35} \times C.SPH$ | – | – | 3.00000 | – | – | 3.00000 |
| 36 | C.SPH → C.S1P | $\frac{Vm_{36}\times C.SPH}{Km_{36}+C.SPH}$ | 0.00340 | 0.00360 | – | 0.00340 | 0.00100 | – |
| 37 | C.S1P → ER.S1P | $k_{37} \times C.S1P - r_{37} \times ER.S1P$ | – | – | 4.50000 | – | – | 4.50000 |
| 38 | ER.CER → M.CER | $k_{38} \times ER.CER - r_{38} \times M.CER$ | – | – | 1.00000 | – | – | 1.00000 |
| 39 | ER.CER → GA.CER | $k_{39} \times ER.CER - r_{39} \times GA.CER$ | – | – | 5.00000 | – | – | 1.00000 |
| 40 | ER.CER → GA.CER | $k_{40} \times ER.CER$ | – | – | 0.08000 | – | – | 0.02000 |
| 41 | GACF.CER → GACF.GluCER | $\frac{Vm_{41}\times GACF.CER}{Km_{41}+GACF.CER}$ | 0.04000 | 0.01000 | – | 0.40000 | 0.01000 | – |
| 42 | GACF.GluCER → GA.GluCER | $k_{42} \times GACF.GluCER$ | – | – | 1.00000 | – | – | 1.00000 |
| 43 | GA.GluCER → GA.LacCER | $\frac{Vm_{43}\times GA.GluCER}{Km_{43}+GA.GluCER}$ | 0.00300 | 0.00100 | – | 0.00300 | 0.00100 | – |
| 44 | GA.LacCER → GA.GSL | $\frac{Vm_{44}\times GA.LacCER}{Km_{44}+GA.LacCER}$ | 0.00300 | 0.00100 | – | 0.00300 | 0.00100 | – |
| 45 | GA.GSL → OM.GSL | $k_{45} \times GA.GSL$ | – | – | 0.20000 | – | – | 0.20000 |
| 46 | OM.GSL → L.GSL | $k_{46} \times OM.GSL$ | – | – | 0.03000 | – | – | 0.03000 |
| 47 | L.GSL → L.CER | $\frac{Vm_{47}\times GA.LacCER}{Km_{44}+GA.LacCER}$ | 0.01900 | 0.00610 | – | 0.01900 | 0.00610 | – |
| 48 | OM.SM → L.SM | $k_{48} \times OM.SM$ | – | – | 0.00200 | – | – | 0.01100 |
| 49 | L.SM → L.CER | $\frac{Vm_{49}\times L.SM}{(Km_{49}+L.SM)\times(1+GA.C1P/Ki_{C1P})\times(1+C.S1P/Ki_{S1P})}$ | 0.04550 | 0.00183 | – | 0.04550 | 0.01000 | – |
| 50 | L.CER → L.SPH | $\frac{Vm_{50}\times L.CER}{Km_{50}+L.CER}$ | 0.14900 | 0.10000 | – | 0.14900 | 0.00669 | – |
| 51 | L.SPH → C.SPH | $k_{51} \times L.SPH$ | – | – | 1.00000 | – | – | 1.00000 |
| 52 | GA.SPH → C.SPH | $k_{52} \times GA.SPH - r_{52} \times C.SPH$ | – | – | 0.20000 | – | – | 0.20000 |
| 53 | GA.SPH → GA.S1P | $\frac{Vm_{53}\times GA.SPH}{Km_{53}+GA.SPH}$ | 0.00506 | 0.13000 | – | 0.00506 | 0.08000 | – |
| 54 | GA.S1P → GA.SPH | $\frac{Vm_{54}\times GA.S1P}{Km_{54}+GA.S1P}$ | 0.03600 | 2.00000 | – | 0.03600 | 2.00000 | – |
| 55 | GA.CER → GA.SM | $\frac{Vm_{55}\times GA.CER}{Km_{55}+GA.CER}$ | 0.02000 | 0.30000 | – | 0.02000 | 0.30000 | – |
| 56 | GA.SM → OM.SM | $k_{56} \times GA.SM$ | – | – | 0.01500 | – | – | 0.04500 |
| 57 | GA.SM → GA.CER | $\frac{Vm_{57}\times GA.SM}{Km_{57}+GA.SM}$ | 0.14800 | 0.05000 | – | 0.14800 | 0.07000 | – |
| 58 | GA.CER → GA.SPH | $\frac{Vm_{58}\times GA.CER}{Km_{58}+GA.CER}$ | 0.08100 | 0.80000 | – | 0.08100 | 0.52900 | – |
| 59 | GA.CER → GA.C1P | $\frac{Vm_{59}\times GA.CER}{Km_{59}+GA.CER}$ | 0.10700 | 2.00000 | – | 0.10700 | 5.00000 | – |
| 60 | GA.C1P → GA.CER | $\frac{Vm_{60}\times GA.C1P}{Km_{60}+GA.C1P}$ | 0.03600 | 0.75000 | – | 0.03600 | 0.75000 | – |
| 61 | GA.C1P → OM.C1P | $k_{61} \times GA.C1P$ | – | – | 2.50000 | – | – | 2.50000 |
| 62 | OM.C1P → OM.CER | $\frac{Vm_{62}\times OM.C1P}{Km_{62}+OM.C1P}$ | 0.03600 | 0.78000 | – | 0.03600 | 0.78000 | – |
| 63 | OM.CER → OM.C1P | $\frac{Vm_{63}\times OM.CER}{Km_{63}+OM.CER}$ | 0.10700 | 2.10000 | – | 0.10700 | 0.50000 | – |
| 64 | → OM.SM | $k_{64}$ | – | – | 0 | – | – | 0 |
| 65 | → OM.C1P | $k_{65} - r_{65} \times OM.C1P$ | – | – | 0 | – | – | 0 |
| 66 | → OM.CER | $k_{66}$ | – | – | 0 | – | – | 0 |
| 67 | → OM.SPH | $k_{67} - r_{67} \times OM.SPH$ | – | – | 0 | – | – | 0 |
| 68 | → OM.S1P | $k_{68} - r_{68} \times OM.S1P$ | – | – | 0 | – | – | 0 |
| 69 | OM.S1P → OM.SPH | $\frac{Vm_{69}\times OM.S1P}{Km_{69}+OM.S1P}$ | 0.03600 | 0.43000 | – | 0.03600 | 0.43000 | – |

Table S3: System of ordinary differential equations of the sphingolipid network.

$$\frac{d[GA.C1P]}{dt} = \frac{Vm_{59} \times [GA.CER]}{Km_{59} + [GA.CER]} - \frac{Vm_{60} \times [GA.C1P]}{Km_{60} + [GA.C1P]} - k_{61} \times [GA.C1P] \quad (1)$$

$$\frac{d[OM.C1P]}{dt} = k_{61} \times [GA.C1P] - \frac{Vm_{62} \times [OM.C1P]}{Km_{62} + [OM.C1P]} + \frac{Vm_{63} \times [OM.CER]}{Km_{63} + [OM.CER]} + k_{65} - r_{65} \times [OM.C1P] \quad (2)$$

$$\frac{d[ER.CER]}{dt} = \frac{Vm_1}{\left(\frac{1+[C.S1P]}{Ki_{S1P}} \times \frac{1+[GA.C1P]}{Ki_{C1P}}\right)} - \frac{Vm_2 \times [ER.CER]}{Km_2 + [ER.CER]} + \frac{Vm_3 \times [ER.SPH]}{Km_3 + [ER.SPH]}$$
$$- (k_7 \times [ER.CER] - r_7 \times [N.CER]) - (k_{38} \times [ER.CER] - r_{38} \times [M.CER]) - (k_{39} \times [ER.CER]) - (k_{40} \times [ER.CER]) \quad (3)$$

$$\frac{d[IM.CER]}{dt} = \frac{Vm_{28} \times [IM.SM]}{Km_{28} + [IM.SM]} - \frac{Vm_{29} \times [IM.CER]}{Km_{29} + [IM.CER]} - (k_{31} \times [IM.CER] - r_{31} \times [OM.CER]) \quad (4)$$

$$\frac{d[L.CER]}{dt} = \frac{Vm_{47} \times [L.GSL]}{Km_{47} + [L.GSL]} + \frac{Vm_{49} \times [L.SM]}{(Km_{49} + [L.SM]) \times (1 + [GA.C1P]/Ki_{C1P}) \times (1 + [C.S1P]/Ki_{S1P})} - \frac{Vm_{50} \times [L.CER]}{Km_{50} + [L.CER]} \quad (5)$$

$$\frac{d[GA.CER]}{dt} = (k_{39} \times [ER.CER]) - \frac{Vm_{55} \times [GA.CER]}{Km_{55} + [GA.CER]} + \frac{Vm_{57} \times [GA.SM]}{Km_{57} + [GA.SM]} - \frac{Vm_{58} \times [GA.CER]}{Km_{58} + [GA.CER]} - \frac{Vm_{59} \times [GA.CER]}{Km_{59} + [GA.CER]} + \frac{Vm_{60} \times [GA.C1P]}{Km_{60} + [GA.C1P]} \quad (6)$$

$$\frac{d[GACF.CER]}{dt} = (k_{40} \times [ER.CER]) - \frac{Vm_{41} \times [GACF.CER]}{Km_{41} + [GACF.CER]} \quad (7)$$

$$\frac{d[M.CER]}{dt} = \frac{Vm_{19} \times [M.SPH]}{Km_{19} + [M.SPH]} - \frac{Vm_{20} \times [M.CER]}{Km_{20} + [M.CER]} + (k_{38} \times [ER.CER] - r_{38} \times [M.CER]) \quad (8)$$

$$\frac{d[N.CER]}{dt} = (k_7 \times [ER.CER] - r_7 \times [N.CER]) - \frac{Vm_8 \times [N.CER]}{Km_8 + [N.CER]} + \frac{Vm_9 \times [N.SM]}{Km_9 + [N.SM]} - \frac{Vm_{10} \times [N.CER]}{Km_{10} + [N.CER]} \quad (9)$$

$$\frac{d[OM.CER]}{dt} = \frac{Vm_{25} \times [OM.SM]}{Km_{25} + [OM.SM]} - \frac{Vm_{26} \times [OM.CER]}{Km_{26} + [OM.CER]} + (k_{31} \times [IM.CER] - r_{31} \times [OM.CER]) + \frac{Vm_{62} \times [OM.C1P]}{Km_{62} + [OM.C1P]} - \frac{Vm_{63} \times [OM.CER]}{Km_{63} + [OM.CER]} + (k_{66}) \quad (10)$$

$$\frac{d[GA.GluCER]}{dt} = (k_{42} \times [GACF.GluCER]) - \frac{Vm_{43} \times [GA.GluCER]}{Km_{43} + [GA.GluCER]} \quad (11)$$

$$\frac{d[GACF.GluCER]}{dt} = \frac{Vm_{41} \times [GACF.CER]}{Km_{41} + [GACF.CER]} - (k_{42} \times [GACF.GluCER]) \quad (12)$$

$$\frac{d[L.GSL]}{dt} = (k_{46} \times [OM.GSL]) - \frac{Vm_{47} \times [L.GSL]}{Km_{47} + [L.GSL]} \quad (13)$$

$$\frac{d[GA.GSL]}{dt} = \frac{Vm_{44} \times [GA.LacCER]}{Km_{44} + [GA.LacCER]} - (k_{45} \times [GA.GSL]) \quad (14)$$

$$\frac{d[OM.GSL]}{dt} = (k_{45} \times [GA.GSL]) - (k_{46} \times [OM.GSL]) \quad (15)$$

$$\frac{d[GA.LacCER]}{dt} = \frac{Vm_{43} \times [GA.GluCER]}{Km_{43} + [GA.GluCER]} - \frac{Vm_{44} \times [GA.LacCER]}{Km_{44} + [GA.LacCER]} \quad (16)$$

$$\frac{d[C.S1P]}{dt} = (k_{15} \times [N.S1P] - r_{15} \times [C.S1P]) - (k_{16} \times [C.S1P] - r_{16} \times [M.S1P]) + (k_{34} \times [IM.S1P] - r_{34} \times [C.S1P]) + \frac{Vm_{36} \times [C.SPH]}{Km_{36} + [C.SPH]}$$
$$- (k_{37} \times [C.S1P] - r_{37} \times [ER.S1P]) \quad (17)$$

$$\frac{d[ER.S1P]}{dt} = \frac{Vm_4 \times [ER.SPH]}{Km_4 + [ER.SPH]} - \frac{Vm_5 \times [ER.S1P]}{Km_5 + [ER.S1P]} - \frac{Vm_6 \times [ER.S1P]}{Km_6 + [ER.S1P]} + (k_{37} \times [C.S1P] - r_{37} \times [ER.S1P]) \quad (18)$$

$$\frac{d[IM.S1P]}{dt} = \frac{Vm_{30} \times [IM.SPH]}{Km_{30} + [IM.SPH]} - (k_{33} \times [IM.S1P]) - (k_{34} \times [IM.S1P] - r_{34} \times [C.S1P]) \quad (19)$$

$$\frac{d[GA.S1P]}{dt} = \frac{Vm_{53} \times [GA.SPH]}{Km_{53} + [GA.SPH]} - \frac{Vm_{54} \times [GA.S1P]}{Km_{54} + [GA.S1P]} \quad (20)$$

$$\frac{d[M.S1P]}{dt} = (k_{16} \times [C.S1P] - r_{16} \times [M.S1P]) - \frac{Vm_{17} \times [M.S1P]}{Km_{17} + [M.S1P]} + \frac{Vm_{18} \times [M.SPH]}{Km_{18} + [M.SPH]} \quad (21)$$

$$\frac{d[N.S1P]}{dt} = \frac{Vm_{11} \times [N.SPH]}{Km_{11} + [N.SPH]} - \frac{Vm_{12} \times [N.S1P]}{Km_{12} + [N.S1P]} - (k_{15} \times [N.S1P] - r_{15} \times [C.S1P]) \quad (22)$$

$$\frac{d[OM.S1P]}{dt} = \frac{Vm_{27} \times [OM.SPH]}{Km_{27} + [OM.SPH]} + (k_{33} \times [IM.S1P]) + (k_{68} - r_{68} \times [OM.S1P]) - \frac{Vm_{69} \times [OM.S1P]}{Km_{69} + [OM.S1P]} \quad (23)$$

$$\frac{d[ER.SM]}{dt} = (k_{13} \times [N.SM] - r_{13} \times [ER.SM]) - (k_{23} \times [ER.SM] - r_{23} \times [IM.SM]) - (k_{24} \times [ER.SM] - r_{24} \times [OM.SM]) \quad (24)$$

$$\frac{d[IM.SM]}{dt} = (k_{23} \times [ER.SM] - r_{23} \times [IM.SM]) - \frac{Vm_{28} \times [IM.SM]}{Km_{28} + [IM.SM]} \quad (25)$$

$$\frac{d[L.SM]}{dt} = (k_{48} \times [OM.SM]) - \frac{Vm_{49} \times [L.SM]}{(Km_{49} + [L.SM]) \times (1 + [GA.C1P]/Ki_{C1P}) \times (1 + [C.S1P]/Ki_{S1P})} \quad (26)$$

$$\frac{d[GA.SM]}{dt} = \frac{Vm_{55} \times [GA.CER]}{Km_{55} + [GA.CER]} - (k_{56} \times [GA.SM]) - \frac{Vm_{57} \times [GA.SM]}{Km_{57} + [GA.SM]} \quad (27)$$

$$\frac{d[N.SM]}{dt} = \frac{Vm_8 \times [N.CER]}{Km_8 + [N.CER]} - \frac{Vm_9 \times [N.SM]}{Km_9 + [N.SM]} - (k_{13} \times [N.SM] - r_{13} \times [ER.SM]) \quad (28)$$

$$\frac{d[OM.SM]}{dt} = (k_{24} \times [ER.SM] - r_{24} \times [OM.SM]) - \frac{Vm_{25} \times [OM.SM]}{Km_{25} + [OM.SM]} - (k_{48} \times [OM.SM]) + (k_{56} \times [GA.SM]) + (k_{64}) \quad (29)$$

$$\frac{d[C.SPH]}{dt} = (k_{14} \times [N.SPH] - r_{14} \times [C.SPH]) + (k_{21} \times [M.SPH] - r_{21} \times [C.SPH]) - (k_{22} \times [C.SPH] - r_{22} \times [ER.SPH])$$
$$+ (k_{35} \times [IM.SPH] - r_{35} \times [C.SPH]) - \frac{Vm_{36} \times [C.SPH]}{Km_{36} + [C.SPH]} + (k_{51} \times [L.SPH]) - (k_{52} \times [C.SPH] - r_{52} \times [GA.SPH]) \quad (30)$$

$$\frac{d[ER.SPH]}{dt} = \frac{Vm_2 \times [ER.CER]}{Km_2 + [ER.CER]} - \frac{Vm_3 \times [ER.SPH]}{Km_3 + [ER.SPH]} - \frac{Vm_4 \times [ER.SPH]}{Km_4 + [ER.SPH]} + \frac{Vm_5 \times [ER.S1P]}{Km_5 + [ER.S1P]} + (k_{22} \times [C.SPH] - r_{22} \times [ER.SPH]) \quad (31)$$

$$\frac{d[IM.SPH]}{dt} = \frac{Vm_{29} \times [IM.CER]}{Km_{29} + [IM.CER]} - \frac{Vm_{30} \times [IM.SPH]}{Km_{30} + [IM.SPH]} - (k_{32} \times [IM.SPH] - r_{32} \times [OM.SPH]) - (k_{35} \times [IM.SPH] - r_{35} \times [C.SPH]) \quad (32)$$

$$\frac{d[L.SPH]}{dt} = \frac{Vm_{50} \times [L.CER]}{Km_{50} + [L.CER]} - (k_{51} \times [L.SPH]) \quad (33)$$

$$\frac{\mathrm{d}[GA.SPH]}{\mathrm{d}t} = (k_{52} \times [C.SPH] - r_{52} \times [GA.SPH]) - \frac{Vm_{53} \times [GA.SPH]}{Km_{53} + [GA.SPH]} + \frac{Vm_{54} \times [GA.S1P]}{Km_{54} + [GA.S1P]} + \frac{Vm_{58} \times [GA.CER]}{Km_{58} + [GA.CER]} \quad (34)$$

$$\frac{\mathrm{d}[M.SPH]}{\mathrm{d}t} = \frac{Vm_{17} \times [M.S1P]}{Km_{17} + [M.S1P]} - \frac{Vm_{18} \times [M.SPH]}{Km_{18} + [M.SPH]} - \frac{Vm_{19} \times [M.SPH]}{Km_{19} + [M.SPH]} + \frac{Vm_{20} \times [M.CER]}{Km_{20} + [M.CER]} - (k_{21} \times [M.SPH] - r_{21} \times [C.SPH]) \quad (35)$$

$$\frac{\mathrm{d}[N.SPH]}{\mathrm{d}t} = \frac{Vm_{10} \times [N.CER]}{Km_{10} + [N.CER]} - \frac{Vm_{11} \times [N.SPH]}{Km_{11} + [N.SPH]} + \frac{Vm_{12} \times [N.S1P]}{Km_{12} + [N.S1P]} - (k_{14} \times [N.SPH] - r_{14} \times [C.SPH]) \quad (36)$$

$$\frac{\mathrm{d}[OM.SPH]}{\mathrm{d}t} = \frac{Vm_{26} \times [OM.CER]}{Km_{26} + [OM.CER]} - \frac{Vm_{27} \times [OM.SPH]}{Km_{27} + [OM.SPH]} + (k_{32} \times [IM.SPH] - r_{32} \times [OM.SPH]) + (k_{67} - r_{67} \times [OM.SPH]) + \frac{Vm_{69} \times [OM.S1P]}{Km_{69} + [OM.S1P]} \quad (37)$$

Table S4: Steady state sphingolipid concentrations for the wildtype and Alzheimer's disease kinetic model.

| # | Class | Wildtype concentration (nmol/mg) | Alzheimer's disease concentration (nmol/mg) |
|---|---|---|---|
| 1 | GA.C1P | 0.001459 | 0.002544 |
| 2 | OM.C1P | 0.002373 | 0.001674 |
| 3 | ER.CER | 0.004556 | 0.015994 |
| 4 | IM.CER | 0.002204 | 0.006029 |
| 5 | L.CER | 0.002455 | 0.191720 |
| 6 | GA.CER | 0.001788 | 0.001209 |
| 7 | GACF.CER | 0.001513 | 0.001322 |
| 8 | M.CER | 0.004094 | 0.007883 |
| 9 | N.CER | 0.001980 | 0.057990 |
| 10 | OM.CER | 0.002322 | 0.006417 |
| 11 | GA.GluCER | 0.001720 | 0.001411 |
| 12 | GACF.GluCER | 0.000364 | 0.000320 |
| 13 | L.GSL | 0.001207 | 0.001051 |
| 14 | GA.GSL | 0.001822 | 0.001599 |
| 15 | OM.GSL | 0.012148 | 0.010663 |
| 16 | GA.LacCER | 0.001720 | 0.001411 |
| 17 | C.S1P | 0.000327 | 0.000154 |
| 18 | ER.S1P | 0.001218 | 0.008694 |
| 19 | IM.S1P | 0.000620 | 0.000438 |
| 20 | GA.S1P | 0.000663 | 0.000218 |
| 21 | M.S1P | 0.001021 | 0.000801 |
| 22 | N.S1P | 0.001135 | 0.000592 |
| 23 | OM.S1P | 0.000645 | 0.000503 |
| 24 | ER.SM | 0.160374 | 0.243261 |
| 25 | IM.SM | 0.132108 | 0.199413 |
| 26 | L.SM | 0.105423 | 0.024589 |
| 27 | GA.SM | 0.123667 | 0.041210 |
| 28 | N.SM | 0.011979 | 0.041789 |
| 29 | OM.SM | 0.628298 | 0.313096 |
| 30 | C.SPH | 0.001162 | 0.001432 |
| 31 | ER.SPH | 0.003909 | 0.006367 |
| 32 | IM.SPH | 0.001796 | 0.003148 |
| 33 | L.SPH | 0.001621 | 0.003764 |
| 34 | GA.SPH | 0.001945 | 0.000896 |
| 35 | M.SPH | 0.003631 | 0.023567 |
| 36 | N.SPH | 0.004038 | 0.004664 |
| 37 | OM.SPH | 0.002293 | 0.003768 |

Table S5: Steady state sphingolipid fluxes for the wildtype and Alzheimer's disease kinetic model.

| # | Reaction | Wildtype flux (nmol/mg/min) | Alzheimer's disease flux (nmol/mg/min) |
|---|---|---|---|
| 1 | → ER.CER | 0 | 0 |
| 2 | ER.CER → ER.SPH | 0.025560 | 0.052766 |
| 3 | ER.SPH → ER.CER | 0.053634 | 0.086157 |
| 4 | ER.SPH → ER.S1P | 0.093591 | 0.570411 |
| 5 | ER.S1P → ER.SPH | 0.095056 | 0.571062 |
| 6 | ER.S1P → ER.PhET + 2THD | 0 | 0 |
| 7 | ER.CER → N.CER | 0.003446 | 0.006996 |
| 8 | N.CER → N.SM | 0.001375 | 0.029677 |
| 9 | N.SM → N.CER | 0.000098 | 0.024906 |
| 10 | N.CER → N.SPH | 0.002169 | 0.002224 |
| 11 | N.SPH → N.S1P | 0.054287 | 0.028918 |
| 12 | N.S1P → N.SPH | 0.053869 | 0.028696 |
| 13 | N.SM → ER.SM | 0.001277 | 0.004771 |
| 14 | N.SPH → C.SPH | 0.001752 | 0.002002 |
| 15 | N.S1P → C.S1P | 0.000418 | 0.000222 |
| 16 | C.S1P → M.S1P | 0.000037 | 0.000003 |
| 17 | M.S1P → M.SPH | 0.077500 | 0.061178 |
| 18 | M.SPH → M.S1P | 0.077463 | 0.061174 |
| 19 | M.SPH → M.CER | 0.059223 | 0.090409 |
| 20 | M.CER → M.SPH | 0.060708 | 0.100491 |
| 21 | M.SPH → C.SPH | 0.001522 | 0.010085 |
| 22 | C.SPH → ER.SPH | 0.026610 | 0.032739 |
| 23 | ER.SM → IM.SM | 0.000943 | 0.001436 |
| 24 | ER.SM → OM.SM | 0.000334 | 0.003336 |
| 25 | OM.SM → OM.CER | 0.000932 | 0.001746 |
| 26 | OM.CER → OM.SPH | 0.004463 | 0.007718 |
| 27 | OM.SPH → OM.S1P | 0.006950 | 0.005487 |
| 28 | IM.SM → IM.CER | 0.000943 | 0.001435 |
| 29 | IM.CER → IM.SPH | 0.001061 | 0.001823 |
| 30 | IM.SPH → IM.S1P | 0.000786 | 0.000575 |
| 31 | OM.CER → IM.CER | 0.000118 | 0.000388 |
| 32 | OM.SPH → IM.SPH | 0.005083 | 0.008156 |
| 33 | IM.S1P → OM.S1P | 0.000620 | 0.000438 |
| 34 | IM.S1P → C.S1P | 0.000166 | 0.000137 |
| 35 | IM.SPH → C.SPH | 0.005358 | 0.009404 |
| 36 | C.SPH → C.S1P | 0.000917 | 0.000296 |
| 37 | C.S1P → ER.S1P | 0.001464 | 0.000652 |
| 38 | ER.CER → M.CER | 0.001485 | 0.010082 |
| 39 | ER.CER → GA.CER | 0.022779 | 0.015994 |
| 40 | ER.CER → GA.CER | 0.000364 | 0.000320 |
| 41 | GACF.CER → GACF.GluCER | 0.000364 | 0.000320 |
| 42 | GACF.GluCER → GA.GluCER | 0.000364 | 0.000320 |
| 43 | GA.GluCER → GA.LacCER | 0.000364 | 0.000320 |
| 44 | GA.LacCER → GA.GSL | 0.000364 | 0.000320 |
| 45 | GA.GSL → OM.GSL | 0.000364 | 0.000320 |
| 46 | OM.GSL → L.GSL | 0.000364 | 0.000320 |
| 47 | L.GSL → L.CER | 0.000364 | 0.000320 |
| 48 | OM.SM → L.SM | 0.001256 | 0.003444 |
| 49 | L.SM → L.CER | 0.001256 | 0.003444 |
| 50 | L.CER → L.SPH | 0.001621 | 0.003764 |
| 51 | L.SPH → C.SPH | 0.001621 | 0.003764 |
| 52 | GA.SPH → C.SPH | 0.017275 | 0.007779 |
| 53 | GA.SPH → GA.S1P | 0.036100 | 0.012037 |
| 54 | GA.S1P → GA.SPH | 0.036100 | 0.012037 |
| 55 | GA.CER → GA.SM | 0.024616 | 0.017100 |
| 56 | GA.SM → OM.SM | 0.001855 | 0.001854 |
| 57 | GA.SM → GA.CER | 0.022761 | 0.015246 |
| 58 | GA.CER → GA.SPH | 0.017275 | 0.007779 |
| 59 | GA.CER → GA.C1P | 0.032867 | 0.055861 |
| 60 | GA.C1P → GA.CER | 0.029218 | 0.049501 |
| 61 | GA.C1P → OM.C1P | 0.003648 | 0.006360 |
| 62 | OM.C1P → OM.CER | 0.048244 | 0.034650 |
| 63 | OM.CER → OM.C1P | 0.044595 | 0.028290 |
| 64 | → OM.SM | 0 | 0 |
| 65 | → OM.C1P | 0 | 0 |
| 66 | → OM.CER | 0 | 0 |
| 67 | → OM.SPH | 0 | 0 |
| 68 | → OM.S1P | 0 | 0 |
| 69 | OM.S1P → OM.SPH | 0.007570 | 0.005926 |

### 4 Sphingolipid network elementary flux modes

Table S6: Description of each EFM in Table S7.

| EFM | Metabolite sequence |
|---|---|
| 1 | GA.SM → GA.CER → GA.SM |
| 2 | ER.CER → GA.CER → GA.SM → OM.SM → OM.CER → IM.CER → IM.SPH → IM.S1P → C.S1P → M.S1P → M.SPH → C.SPH → ER.SPH → ER.CER |
| 3 | ER.CER → ER.SPH → ER.CER |
| 4 | M.SPH → C.SPH → ER.SPH → ER.CER → N.CER → N.SM → ER.SM → OM.SM → OM.CER → OM.SPH → IM.SPH → IM.S1P → C.S1P → M.S1P → M.SPH |
| 5 | M.SPH → M.S1P → M.SPH |
| 6 | N.S1P → N.SPH → N.S1P |
| 7 | N.S1P → C.S1P → ER.S1P → ER.SPH → ER.CER → N.CER → N.SPH → N.S1P |
| 8 | GA.S1P → GA.SPH → GA.S1P |
| 9 | M.SPH → C.SPH → C.S1P → M.S1P → M.SPH |
| 10 | L.SPH → C.SPH → ER.SPH → ER.CER → N.CER → N.SM → ER.SM → OM.SM → L.SM → L.CER → L.SPH |
| 11 | N.SM → N.CER → N.SM |
| 12 | M.SPH → M.CER → M.SPH |
| 13 | M.SPH → C.SPH → ER.SPH → ER.CER → M.CER → M.SPH |
| 14 | C.SPH → ER.SPH → ER.CER → GACF.CER → GACF.GluCER → GA.GluCER → GA.LacCER → GA.GSL → OM.GSL → L.GSL → L.CER → L.SPH → C.SPH |
| 15 | C.SPH → C.S1P → ER.S1P → ER.SPH → ER.CER → GACF.CER → GACF.GluCER → GA.GluCER → GA.LacCER → GA.GSL → OM.GSL → L.GSL → L.CER → L.SPH → C.SPH |
| 16 | IM.CER → IM.SPH → C.SPH → ER.SPH → ER.CER → N.CER → N.SM → ER.SM → IM.SM → IM.CER |
| 17 | OM.SPH → IM.SPH → IM.S1P → OM.S1P → OM.SPH |
| 18 | OM.SPH → OM.S1P → OM.SPH |
| 19 | IM.SM → IM.CER → IM.SPH → C.SPH → C.S1P → ER.S1P → ER.SPH → ER.CER → N.CER → N.SM → ER.SM → IM.SM |
| 20 | GA.SPH → C.SPH → ER.SPH → ER.CER → GA.CER → GA.SPH |
| 21 | IM.SM → IM.CER → IM.SPH → IM.S1P → C.S1P → M.S1P → M.SPH → C.SPH → ER.SPH → ER.CER → N.CER → N.SM → ER.SM → IM.SM |
| 22 | OM.SM → OM.CER → IM.CER → IM.SPH → IM.S1P → C.S1P → ER.S1P → ER.SPH → ER.CER → GA.CER → GA.SM → OM.SM |
| 23 | N.SPH → C.SPH → C.S1P → ER.S1P → ER.SPH → ER.CER → N.CER → N.SPH |
| 24 | IM.SM → IM.CER → IM.SPH → IM.S1P → C.S1P → ER.S1P → ER.SPH → ER.CER → N.CER → N.SM → ER.SM → IM.SM |
| 25 | ER.CER → GA.CER → GA.SM → OM.SM → L.SM → L.CER → L.SPH → C.SPH → ER.SPH → ER.CER |
| 26 | ER.SM → OM.SM → L.SM → L.CER → L.SPH → C.SPH → C.S1P → ER.S1P → ER.SPH → ER.CER → N.CER → N.SM → ER.SM |
| 27 | M.SPH → C.SPH → C.S1P → ER.S1P → ER.SPH → ER.CER → M.CER → M.SPH |
| 28 | OM.SM → OM.CER → OM.SPH → IM.SPH → IM.S1P → C.S1P → ER.S1P → ER.SPH → ER.CER → GA.CER → GA.SM → OM.SM |
| 29 | OM.SM → OM.CER → IM.CER → IM.SPH → IM.S1P → C.S1P → ER.S1P → ER.SPH → ER.CER → N.CER → N.SM → ER.SM → OM.SM |
| 30 | ER.S1P → ER.SPH → ER.S1P |
| 31 | OM.SM → OM.CER → OM.SPH → IM.SPH → IM.S1P → C.S1P → ER.S1P → ER.SPH → ER.CER → N.CER → N.SM → ER.SM → OM.SM |
| 32 | ER.CER → GA.CER → GA.SM → OM.SM → OM.CER → OM.SPH → IM.SPH → C.SPH → ER.SPH → ER.CER |
| 33 | ER.SM → OM.SM → OM.CER → OM.SPH → IM.SPH → C.SPH → C.S1P → ER.S1P → ER.SPH → ER.CER → N.CER → N.SM → ER.SM |
| 34 | N.S1P → C.S1P → M.S1P → M.SPH → C.SPH → ER.SPH → ER.CER → N.CER → N.SPH → N.S1P |
| 35 | N.SPH → C.SPH → ER.SPH → ER.CER → N.CER → N.SPH |
| 36 | GA.CER → GA.C1P → OM.C1P → OM.CER → IM.CER → IM.SPH → C.SPH → C.S1P → ER.S1P → ER.SPH → ER.CER → GA.CER |
| 37 | GA.CER → GA.C1P → OM.C1P → OM.CER → OM.SPH → IM.SPH → C.SPH → C.S1P → ER.S1P → ER.SPH → ER.CER → GA.CER |
| 38 | IM.SPH → C.SPH → ER.SPH → ER.CER → N.CER → N.SM → ER.SM → OM.SM → OM.CER → OM.SPH → IM.SPH |
| 39 | OM.CER → OM.C1P → OM.CER |
| 40 | L.SPH → C.SPH → C.S1P → ER.S1P → ER.SPH → ER.CER → GA.CER → GA.SM → OM.SM → L.SM → L.CER → L.SPH |
| 41 | GA.SPH → C.SPH → C.S1P → ER.S1P → ER.SPH → ER.CER → GA.CER → GA.SPH |
| 42 | ER.CER → GA.CER → GA.SM → OM.SM → OM.CER → OM.SPH → IM.SPH → IM.S1P → C.S1P → M.S1P → M.SPH → C.SPH → ER.SPH → ER.CER |
| 43 | ER.CER → GA.CER → GA.SM → OM.SM → OM.CER → IM.CER → IM.SPH → C.SPH → C.S1P → ER.S1P → ER.SPH → ER.CER |
| 44 | GA.CER → GA.C1P → OM.C1P → OM.CER → IM.CER → IM.SPH → C.SPH → ER.SPH → ER.CER → GA.CER |
| 45 | ER.SM → OM.SM → OM.CER → IM.CER → IM.SPH → C.SPH → ER.SPH → ER.CER → N.CER → N.SM → ER.SM |
| 46 | GA.CER → GA.C1P → OM.C1P → OM.CER → OM.SPH → IM.SPH → C.SPH → ER.SPH → ER.CER → GA.CER |
| 47 | M.SPH → C.SPH → ER.SPH → ER.CER → N.CER → N.SM → ER.SM → OM.SM → OM.CER → IM.CER → IM.SPH → IM.S1P → C.S1P → M.S1P → M.SPH |
| 48 | IM.SPH → C.SPH → C.S1P → ER.S1P → ER.SPH → ER.CER → N.CER → N.SM → ER.SM → OM.SM → OM.CER → IM.CER → IM.SPH |
| 49 | OM.SM → OM.CER → OM.SPH → IM.SPH → C.SPH → C.S1P → ER.S1P → ER.SPH → ER.CER → GA.CER → GA.SM → OM.SM |
| 50 | GA.CER → GA.C1P → OM.C1P → OM.CER → IM.CER → IM.SPH → IM.S1P → C.S1P → ER.S1P → ER.SPH → ER.CER → GA.CER |
| 51 | GA.CER → GA.C1P → OM.C1P → OM.CER → OM.SPH → IM.SPH → IM.S1P → C.S1P → ER.S1P → ER.SPH → ER.CER → GA.CER |
| 52 | GA.CER → GA.C1P → OM.C1P → OM.CER → IM.CER → IM.SPH → IM.S1P → C.S1P → M.S1P → M.SPH → C.SPH → ER.SPH → ER.CER → GA.CER |
| 53 | ER.CER → GA.CER → GA.SM → OM.SM → OM.CER → IM.CER → IM.SPH → C.SPH → ER.SPH → ER.CER |
| 54 | GA.CER → GA.C1P → OM.C1P → OM.CER → OM.SPH → IM.SPH → IM.S1P → C.S1P → M.S1P → M.SPH → C.SPH → ER.SPH → ER.CER → GA.CER |
| 55 | GA.CER → GA.C1P → GA.CER |

### 5 Markov weights for the wildtype and Alzheimer's disease sphingolipid network

Table S7: EFM weights for the Markov method in the wildtype and Alzheimer's disease condition. # of compartments refers to the number of reactions across unique compartments in the sphingolipid network (9 total). EFM length refers to the number of reactions in each EFM.

| EFM | $w_{wt}$ | $w_{ad}$ | $\log_2(w_{ad}/w_{wt})$ | $w_{ad} - w_{wt}$ | # compartments | EFM length |
|---|---|---|---|---|---|---|
| 1 | $2.276 \times 10^{-2}$ | $1.525 \times 10^{-2}$ | $-5.781 \times 10^{-1}$ | $-7.515 \times 10^{-3}$ | 1 | 2 |
| 2 | $1.449 \times 10^{-8}$ | $2.176 \times 10^{-9}$ | $-2.736$ | $-1.232 \times 10^{-8}$ | 6 | 13 |
| 3 | $2.556 \times 10^{-2}$ | $5.277 \times 10^{-2}$ | $1.046$ | $2.721 \times 10^{-2}$ | 1 | 2 |
| 4 | $9.882 \times 10^{-8}$ | $7.779 \times 10^{-8}$ | $-3.452 \times 10^{-1}$ | $-2.103 \times 10^{-8}$ | 6 | 14 |
| 5 | $7.746 \times 10^{-2}$ | $6.117 \times 10^{-2}$ | $-3.406 \times 10^{-1}$ | $-1.629 \times 10^{-2}$ | 1 | 2 |
| 6 | $5.387 \times 10^{-2}$ | $2.870 \times 10^{-2}$ | $-9.086 \times 10^{-1}$ | $-2.517 \times 10^{-2}$ | 1 | 2 |
| 7 | $4.080 \times 10^{-4}$ | $2.208 \times 10^{-4}$ | $-8.855 \times 10^{-1}$ | $-1.871 \times 10^{-4}$ | 3 | 7 |
| 8 | $3.610 \times 10^{-2}$ | $1.204 \times 10^{-2}$ | $-1.585$ | $-2.406 \times 10^{-2}$ | 1 | 2 |
| 9 | $2.293 \times 10^{-5}$ | $1.535 \times 10^{-6}$ | $-3.901$ | $-2.139 \times 10^{-5}$ | 2 | 4 |
| 10 | $1.855 \times 10^{-4}$ | $2.194 \times 10^{-3}$ | $3.564$ | $2.008 \times 10^{-3}$ | 5 | 10 |
| 11 | $9.810 \times 10^{-5}$ | $2.491 \times 10^{-2}$ | $7.988$ | $2.481 \times 10^{-2}$ | 1 | 2 |
| 12 | $5.922 \times 10^{-2}$ | $9.041 \times 10^{-2}$ | $6.103 \times 10^{-1}$ | $3.119 \times 10^{-2}$ | 1 | 2 |
| 13 | $1.437 \times 10^{-3}$ | $9.992 \times 10^{-3}$ | $2.798$ | $8.555 \times 10^{-3}$ | 3 | 5 |
| 14 | $3.526 \times 10^{-4}$ | $3.170 \times 10^{-4}$ | $-1.535 \times 10^{-1}$ | $-3.559 \times 10^{-5}$ | 6 | 12 |
| 15 | $1.185 \times 10^{-5}$ | $2.855 \times 10^{-6}$ | $-2.054$ | $-8.999 \times 10^{-6}$ | 6 | 14 |
| 16 | $8.850 \times 10^{-4}$ | $1.402 \times 10^{-3}$ | $6.640 \times 10^{-1}$ | $5.173 \times 10^{-4}$ | 4 | 9 |
| 17 | $6.198 \times 10^{-4}$ | $4.385 \times 10^{-4}$ | $-4.991 \times 10^{-1}$ | $-1.813 \times 10^{-4}$ | 2 | 4 |
| 18 | $6.950 \times 10^{-3}$ | $5.487 \times 10^{-3}$ | $-3.409 \times 10^{-1}$ | $-1.463 \times 10^{-3}$ | 1 | 2 |
| 19 | $2.975 \times 10^{-5}$ | $1.263 \times 10^{-5}$ | $-1.236$ | $-1.712 \times 10^{-5}$ | 4 | 11 |
| 20 | $1.671 \times 10^{-2}$ | $7.710 \times 10^{-3}$ | $-1.116$ | $-9.004 \times 10^{-3}$ | 3 | 5 |
| 21 | $6.725 \times 10^{-7}$ | $1.045 \times 10^{-7}$ | $-2.686$ | $-5.680 \times 10^{-7}$ | 5 | 13 |
| 22 | $5.972 \times 10^{-7}$ | $4.263 \times 10^{-7}$ | $-4.863 \times 10^{-1}$ | $-1.709 \times 10^{-7}$ | 5 | 11 |
| 23 | $5.697 \times 10^{-5}$ | $1.787 \times 10^{-5}$ | $-1.672$ | $-3.909 \times 10^{-5}$ | 3 | 7 |
| 24 | $2.771 \times 10^{-5}$ | $2.047 \times 10^{-5}$ | $-4.365 \times 10^{-1}$ | $-7.234 \times 10^{-6}$ | 4 | 11 |
| 25 | $1.030 \times 10^{-3}$ | $1.220 \times 10^{-3}$ | $2.434 \times 10^{-1}$ | $1.893 \times 10^{-4}$ | 5 | 9 |
| 26 | $6.234 \times 10^{-6}$ | $1.975 \times 10^{-5}$ | $1.664$ | $1.352 \times 10^{-5}$ | 5 | 12 |
| 27 | $4.830 \times 10^{-5}$ | $8.997 \times 10^{-5}$ | $8.975 \times 10^{-1}$ | $4.167 \times 10^{-5}$ | 3 | 7 |
| 28 | $2.262 \times 10^{-5}$ | $8.472 \times 10^{-6}$ | $-1.417$ | $-1.414 \times 10^{-5}$ | 5 | 11 |
| 29 | $1.075 \times 10^{-7}$ | $7.668 \times 10^{-7}$ | $2.835$ | $6.593 \times 10^{-7}$ | 5 | 12 |
| 30 | $9.359 \times 10^{-2}$ | $5.704 \times 10^{-1}$ | $2.608$ | $4.768 \times 10^{-1}$ | 1 | 2 |
| 31 | $4.071 \times 10^{-6}$ | $1.524 \times 10^{-5}$ | $1.904$ | $1.117 \times 10^{-5}$ | 5 | 12 |
| 32 | $7.224 \times 10^{-4}$ | $5.803 \times 10^{-4}$ | $-3.160 \times 10^{-1}$ | $-1.421 \times 10^{-4}$ | 5 | 9 |
| 33 | $4.372 \times 10^{-6}$ | $9.399 \times 10^{-6}$ | $1.104$ | $5.028 \times 10^{-6}$ | 5 | 12 |
| 34 | $9.903 \times 10^{-6}$ | $1.127 \times 10^{-6}$ | $-3.135$ | $-8.775 \times 10^{-6}$ | 4 | 9 |
| 35 | $1.695 \times 10^{-3}$ | $1.985 \times 10^{-3}$ | $2.280 \times 10^{-1}$ | $2.901 \times 10^{-4}$ | 3 | 5 |
| 36 | $2.961 \times 10^{-6}$ | $2.680 \times 10^{-6}$ | $-1.436 \times 10^{-1}$ | $-2.805 \times 10^{-7}$ | 5 | 11 |
| 37 | $1.121 \times 10^{-4}$ | $5.326 \times 10^{-5}$ | $-1.074$ | $-5.886 \times 10^{-5}$ | 5 | 11 |
| 38 | $1.300 \times 10^{-4}$ | $1.044 \times 10^{-3}$ | $3.005$ | $9.138 \times 10^{-4}$ | 5 | 10 |
| 39 | $4.459 \times 10^{-2}$ | $2.829 \times 10^{-2}$ | $-6.566 \times 10^{-1}$ | $-1.631 \times 10^{-2}$ | 1 | 2 |
| 40 | $3.463 \times 10^{-5}$ | $1.098 \times 10^{-5}$ | $-1.657$ | $-2.365 \times 10^{-5}$ | 5 | 11 |
| 41 | $5.619 \times 10^{-4}$ | $6.942 \times 10^{-5}$ | $-3.017$ | $-4.924 \times 10^{-4}$ | 3 | 7 |
| 42 | $5.490 \times 10^{-7}$ | $4.324 \times 10^{-8}$ | $-3.666$ | $-5.057 \times 10^{-7}$ | 6 | 13 |
| 43 | $6.412 \times 10^{-7}$ | $2.629 \times 10^{-7}$ | $-1.286$ | $-3.783 \times 10^{-7}$ | 5 | 11 |
| 44 | $8.807 \times 10^{-5}$ | $2.976 \times 10^{-4}$ | $1.757$ | $2.096 \times 10^{-4}$ | 5 | 9 |
| 45 | $3.434 \times 10^{-6}$ | $5.252 \times 10^{-5}$ | $3.935$ | $4.909 \times 10^{-5}$ | 5 | 10 |
| 46 | $3.335 \times 10^{-3}$ | $5.915 \times 10^{-3}$ | $8.266 \times 10^{-1}$ | $2.580 \times 10^{-3}$ | 5 | 9 |
| 47 | $2.609 \times 10^{-9}$ | $3.914 \times 10^{-9}$ | $5.850 \times 10^{-1}$ | $1.305 \times 10^{-9}$ | 6 | 14 |
| 48 | $1.154 \times 10^{-7}$ | $4.730 \times 10^{-7}$ | $2.035$ | $3.575 \times 10^{-7}$ | 5 | 12 |
| 49 | $2.428 \times 10^{-5}$ | $5.225 \times 10^{-6}$ | $-2.216$ | $-1.906 \times 10^{-5}$ | 5 | 11 |
| 50 | $2.757 \times 10^{-6}$ | $4.345 \times 10^{-6}$ | $6.563 \times 10^{-1}$ | $1.588 \times 10^{-6}$ | 5 | 11 |
| 51 | $1.044 \times 10^{-4}$ | $8.636 \times 10^{-5}$ | $-2.740 \times 10^{-1}$ | $-1.806 \times 10^{-5}$ | 5 | 11 |
| 52 | $6.692 \times 10^{-8}$ | $2.218 \times 10^{-8}$ | $-1.593$ | $-4.474 \times 10^{-8}$ | 6 | 13 |
| 53 | $1.907 \times 10^{-5}$ | $2.920 \times 10^{-5}$ | $6.142 \times 10^{-1}$ | $1.012 \times 10^{-5}$ | 5 | 9 |
| 54 | $2.535 \times 10^{-6}$ | $4.408 \times 10^{-7}$ | $-2.523$ | $-2.094 \times 10^{-6}$ | 6 | 13 |
| 55 | $2.922 \times 10^{-2}$ | $4.950 \times 10^{-2}$ | $7.606 \times 10^{-1}$ | $2.028 \times 10^{-2}$ | 1 | 2 |

### 6 Individual flux reconstruction error across methods

a) b)

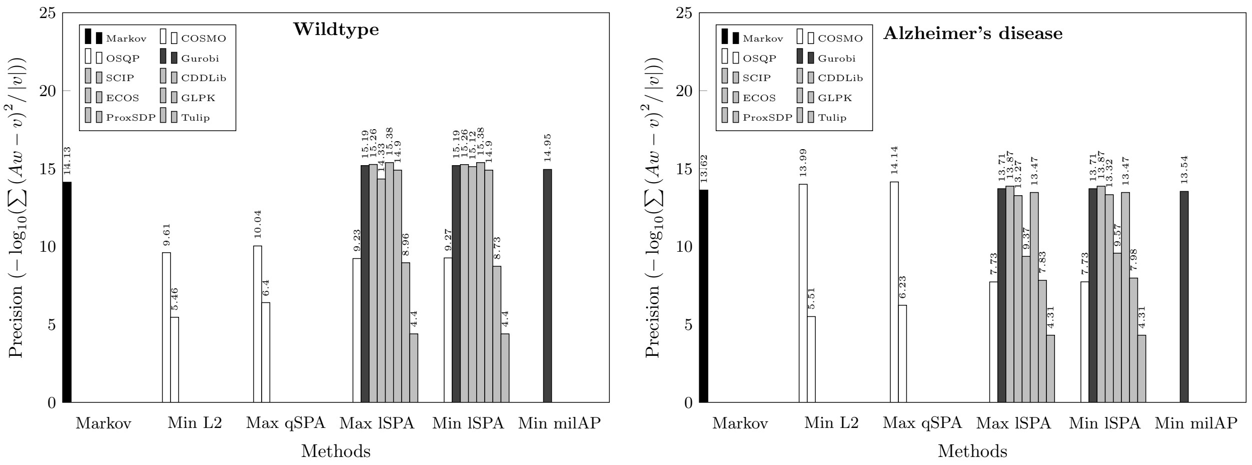

Figure S1: Error between individual fluxes reconstructed from EFM weights from different methods and the observed network fluxes for the a) wildtype sphingolipid network and b) Alzheimer's disease sphingolipid network. $A$ is the binary matrix of EFM weights (rows are reactions, columns are EFMs), $w$ is the set of EFM weights, and $v$ is the vector of observed network fluxes.

### 7 Total flux reconstruction error across methods

a) b)

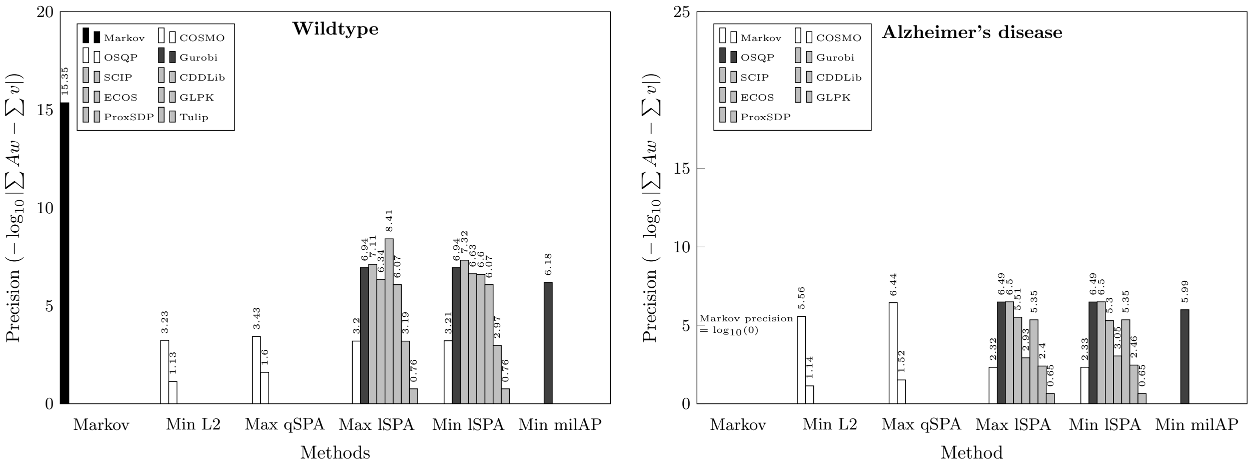

Figure S2: Error between total fluxes reconstructed from EFM weights from different methods and the observed network fluxes for the a) wildtype sphingolipid network and b) Alzheimer's disease sphingolipid network. $A$ is the binary matrix of EFM weights (rows are reactions, columns are EFMs), $w$ is the set of EFM weights, and $v$ is the vector of observed network fluxes.
